# Extreme purifying selection against point mutations in the human genome

**DOI:** 10.1101/2021.08.23.457339

**Authors:** Noah Dukler, Mehreen R. Mughal, Ritika Ramani, Yi-Fei Huang, Adam Siepel

## Abstract

Genome sequencing of tens of thousands of humans has enabled the measurement of large selective effects for mutations to protein-coding genes. Here we describe a new method, called ExtRaINSIGHT, for measuring similar selective effects in noncoding as well as in coding regions of the human genome. ExtRaINSIGHT estimates the prevalance of strong purifying selection, or “ultraselection” (*λ_s_*), as the fractional depletion of rare single-nucleotide variants in target genomic sites relative to matched sites that are putatively free from selection, after controlling for local variation and neighbor-dependence in mutation rate. We show using simulations that *λ_s_* is closely related to the average site-specific selection coefficient against heterozygous point mutations, as predicted at mutation-selection balance. Applying ExtRaINSIGHT to 71,702 whole genome sequences from gnomAD v3, we find strong evidence of ultraselection in evolutionarily ancient miRNAs and neuronal protein-coding genes, as well as at splice sites. By contrast, we find weak evidence in other noncoding RNAs and transcription factor binding sites, and only modest evidence in ultraconserved elements and human accelerated regions. We estimate that ~0.3–0.5% of the human genome is ultraselected, implying ~0.3–0.4 lethal or nearly lethal *de novo* mutations per potential human zygote. Overall, our study sheds new light on the genome-wide distribution of fitness effects for new point mutations by combining deep new sequencing data sets and classical theory from population genetics.

## Introduction

Like a gambler, an evolving species has to pay for the chance to win. As in most games of chance, the majority of “draws” (mutations) result in a loss (decrease in fitness), with an occasional pay-off (adaptive mutation). Thus, in Haldane’s words, loss of fitness owing to deleterious mutation is the “price paid by a species for its capacity for further evolution” [1].

Understanding the impact of new mutations on fitness has been a major focus of evolutionary genetics for nearly a century [1–3], with implications for a wide variety of fundamental problems, ranging from revealing the genetic architecture of complex traits and the effects of mutational load to understanding the emergence of recombination and sex [4, 5]. Nevertheless, characterizing the full distribution of fitness effects (DFE) of new mutations is notoriously difficult. Naturally occurring mutations are rare, often difficult to detect, and have fitness effects that are generally hard to measure. Innovative experimental techniques have been developed to measure of the DFE in model organisms, but these methods have important limitations [4] and, in any case, they cannot be applied to humans, nor to any other organism that cannot be experimentally manipulated and monitored in relatively large numbers.

For these reasons, many recent efforts to characterize the DFE have focused on the study of naturally occurring mutations using statistical modeling, population genetic theory, and DNA sequencing [6–9]. Importantly, however, patterns of genetic variation are strongly influenced by demographic history, so careful demographic modeling is required to isolate the effects of selection. In addition, most available population panels—consisting of hundreds to a few thousand individuals—are informative about only a relatively narrow slice of the DFE. For example, in humans strong purifying selection (such that *s* > ~1%) will tend to hold variants below a detectable frequency in these panels, whereas weak purifying selection (such that *s* < ~10^-4^) will be indistinguishable from random genetic drift [10, 11]. Thus, only in approximately the range 10^-4^ < *s* < 10^-2^ can purifying selection be accurately measured.

Recently, exome or whole-genome sequence data has become available for tens of thousands of individuals [12, 13], allowing quite rare variants (with relative frequencies < 10^-3^) to be identified with reasonable confidence. These data have enabled the application of statistical methods that can measure high levels of purifying selection against predicted loss-of-function (pLoF) mutations for protein-coding genes [11–16]. While such measures are correlated with dominance effects (e.g., [12, 13]), the frequency of rare pLoF variants is strictly informative only about the strength of selection against hetereozygous mutations, *s*_het_ [17]. When purifying selection is strong and near-complete recessivity can be excluded, mutation-selection balance is expected to hold with an equilibrium frequency for a rare variant of 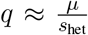, where *μ* is the deleterious mutation rate [1, 17]. Cassa et al. [11] (see also [18]) have shown with extensive simulations that this relationship holds quite well for pLoF variants in the ExAC exome data [12] down to *s*_het_ ≈ 0.01. Importantly, estimation of *s*_het_ based on mutation-selection balance is independent of demography because, in this regime, mutant alleles persist in the population for at most a few generations and genetic drift makes a negligible contribution to their allele frequencies. Therefore, in addition to permitting estimation of larger selection coefficients than other statistical methods, this approach requires no demographic modeling.

In this article, we extend and generalize these ideas for application to the entire genome, including noncoding regions, in a new method called Extremely Rare INSIGHT (ExtRaINSIGHT). Similar to our previous Inference of Natural Selection from Interspersed Genomically coHerent elemenTs (INSIGHT) method [19, 20], ExtRaINSIGHT can be used to measure the influence of natural selection on any designated set of genomic sequences, by contrasting patterns of variation in a designated set of “target” sequences with those in matched sequences that are putatively neutrally evolving. However, ExtRaINSIGHT focuses on rare variants only, in order to obtain a measure that reflects particularly large selective effects— that is, purifying selection sufficiently strong that new point mutations are lethal or nearly lethal (hereafter, “nearly lethal”), and therefore do not appear even as rare variants in a panel of tens of thousands of individuals. As shorthand, we refer to such selection as “ultraselection.” We apply ExtRaINSIGHT to more than 70,000 whole genome sequences from the Genome Aggregation Database (gnomAD) project (https://gnomad.broadinstitute.org/) [13] and perform a comprehensive analysis of ultraselection in the human genome, considering both coding and noncoding elements. Our findings reveal both similarities and striking differences in measures of ultraselection and weaker purifying selection, shed light on the rate of nearly lethal mutations in humans, and highlight challenges in accurately modeling mutation rates in upstream regions of genes.

## Results

### Overview of ExtRaINSIGHT

ExtRaINSIGHT measures the fractional reduction in the incidence of rare variants in a target set of sites relative to nearby sites that are putatively free from (direct) natural selection. In this way, it is analogous to classical strategies for measuring selection in protein-coding genes [21–23], as well as to newer methods that compare target sets of noncoding elements with suitable background sequences [20, 24–26]. The focus on rare variants (here, variants with minor allele frequencies of < 0.1%), however, enables the method to focus in particular on point mutations of large selective effect.

The main challenge in this approach stems from the high sensitivity of relative rates of rare variants to variation in mutation rate. To address this problem, we follow refs. [12, 15] in building a mutational model that accounts for both sequence context and regional variation in mutation rate. In our case, we condition the rate of each type of nucleotide substitution on the identity of the three flanking nucleotides on each side. In addition, following our earlier work [19, 20], we use a local control for overall mutation rate based on nearby sites identified as likely to be neutrally evolving. We also consider G+C content, sequencing coverage, and CpG islands as covariates (see **Methods**). With this strategy, we are able to predict with high accuracy the probability that a rare variant will occur at each site (**Supplemental Fig. S1)**.

In the absence of natural selection, we assume a Bernoulli sampling model for the presence (probability *P_i_*) or absence (probably 1 – *P_i_*) of a rare variant at each site *i*, where *P_i_* reflects the local sequence context and overall rate of mutation. We ignore sites at which common variants occur (similar to [12, 15]). We then assume that natural selection has the effect of imposing a fractional reduction on the rate at which rare variants occur. To a first approximation, we maximize the following likelihood function,

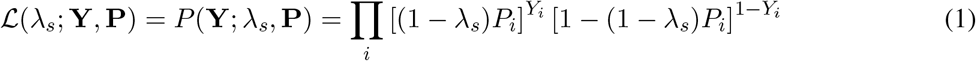

where *Y_i_* is an indicator variable for the presence of a rare variant at position i in the sample, *λ_s_* is a scale factor capturing a depletion of rare genetic variation, **Y** = {*Y_i_*}, **P** = {*P_i_*}, and the product excludes sites having common variants. In this way, we obtain a maximum-likelihood estimate (MLE) of *λ_s_* conditional on pre-estimated values *P_i_*. (In practice, we use a slighly more complicated likelihood function that distinguishes among the possible alternative alleles at each site; see **Methods** for complete details.)

When *λ_s_* falls between 0 and 1 it can be interpreted as a measure of the prevalence of ultraselection. In this case, *λ_s_* can be thought of as the fraction of sites intolerant to heterozygous mutants, although in practice, some sites may be more, and some sites less, intolerant. Notice, however, that *λ_s_* can also take values < 0 if rare variants occur at a higher-than-expected rate in the target set of sites. As we discuss below, we do observe a systematic tendency for *λ_s_* to take negative values in particular classes of sites, likely reflecting the difficulty of precisely specifying the mutational model at these sites. Across most of the genome, however, estimates of *λ_s_* fall between 0 and 1 and show general qualitative agreement with other measures of purifying selection.

Notably, in the case of strong selection against heterozygotes and mutation-selection balance (as detailed by [11, 17]), a relatively simple relationship can be established between *λ_s_* and the site-specific selection coefficient against heterozygous mutations, *s*_het_:

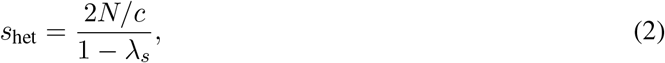

where *N* is the number of diploid individuals sampled and *c* is the (constant for a given data set) ratio of the rate of presence of rare variants in the sample (*P_i_*) to the per-generation mutation rate (see **Methods** and **Supplemental Fig. S2**).

Following ref. [18], we simulated data sets under a realistic human demographic model with various values of *s*_het_ and estimated *λ_s_* from each one using ExtRaINSIGHT. We found that equation 2 led to fairly accurate estimates of the true value down to about *s*_het_ = 0.03, and somewhat elevated but still useful estimates down to about *s*_het_ = 0.013 (**Supplemental Fig. S3**). Therefore, throughout this article, we use equation 2 to estimate *s*_het_ when *λ_s_* > 0.18, approximately the threshold corresponding to *s*_het_ = 0.013 for our data set. Notably, our simulation study did indicate that variation across sites in *s*_het_ leads to some underestimation of the true average value, but even in this setting equation 2 remains useful as an approximate guide (see **Methods** and **Supplemental Fig. S3**).

### Ultraselection in and around protein-coding genes

We applied ExtRaINSIGHT to 19,955 protein-coding genes from GENCODE v. 38 [27] as well as to a variety of proximal coding-associated sequences, including 5’ and 3’ untranslated regions (UTRs), promoters, and splice sites (**Figure 1**). For comparison, we applied INSIGHT to the same sets of elements. As expected, we obtained considerably higher estimates of *λ_s_* at 0-fold degenerate (0d) sites in coding sequences, at which each possible mutation results in an amino-acid change (*λ_s_* = 0.22), than at 4-fold degenerate (4d) sites, at which every mutation is synonymous (*λ_s_* = –0.008). The corresponding INSIGHT-based estimates of *ρ* were 0.80 and 0.39, respectively. Together, we can interpret these estimates as indicating that 22% of 0d sites are ultraselected, meaning that any mutation at these sites would be nearly lethal, and another 80 – 22 = 58% are under weaker purifying selection—although the ExtRaINSIGHT and INSIGHT estimates are not precisely comparable in all respects (see **Discussion**). Our estimate of *λ_s_* for 0d sites corresponds to a selection coefficient of *s*_het_ ≈ 0.014, assuming mutation-selection balance. Notably, this estimate is substantially larger than previous estimates for amino-acid replacing mutations based on the site-frequency-spectrum from smaller samples, probably in part because those methods are less sensitive to strong purifying selection (see **Discussion**). By contrast, at 4d sites, ultraselection is estimated to be completely absent, but 39% of 4d sites experience weak purifying selection (see [9] for an estimate of 26% for synonymous sites). Overall, about 15% of coding sites (CDS) experience ultraselection (*λ_s_* = 0.15) and another 47% experience weaker selection (*ρ* = 0.62).

**Figure 1:**
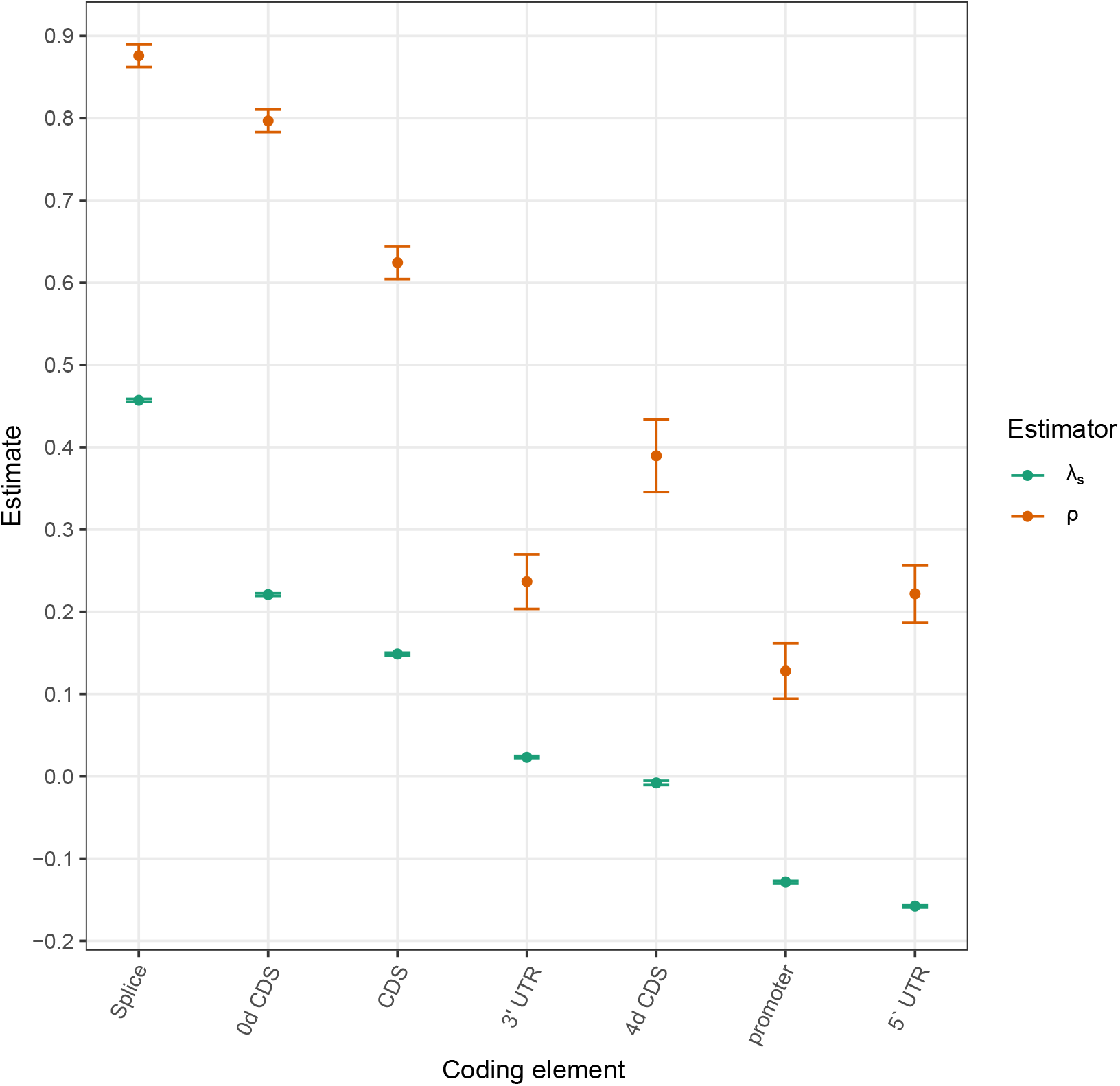
Measures of purifying selection at coding and coding-proximal genomic elements. Estimates are shown for both ExtRaINSIGHT (*λ_s_*) and INSIGHT (*ρ*). Error bars indicate one standard error (see **Methods**).

Among coding-related sites, the strongest selection, by far, occurred in splice sites (see also [28]), where almost half of sites were subject to ultraselection (*λ_s_* = 0.46; corresponding to *s*_het_ ≈ 0.020), with another 42% subject to weaker selection (*ρ* = 0.88). By contrast, 3’ UTRs showed little evidence of ultraselection (*λ_s_* = 0.023) despite considerable evidence of weaker selection (*ρ* = 0.24). Interestingly, we observed a persistent tendency for negative estimates of *λ_s_* at regions near the 5’ ends of genes, at both 5’ UTRs and promoter regions, despite non-neglible estimates of *ρ* (0.22 and 0.13, respectively). As we discuss in a later section, these estimates appear to be a consequence of unusual mutational patterns in these regions that are difficult to accommodate using even our regional and neighbor-dependent mutation model.

To see whether ExtRaINSIGHT was capable of distinguishing among protein-coding sequences experiencing different levels of selection against heterozygous loss-of-function (LoF) variants, we compared it with the recently introduced loss-of-function observed/expected upper bound fraction (LOEUF) measure [13]. LOEUF is similarly based on rare variants but differs from ExtRaINSIGHT in that it is computed separately for each gene by pooling together all mutations predicted to result in loss-of-function of that gene (including nonsense mutations, mutations that disrupt splice sites, and frameshift mutations). In contrast to *λ_s_* and *ρ*, lower LOEUF scores are associated with stronger depletions of LoF variants and increased constraint, and higher LOEUF scores are associated with weaker depletions and reduced constraint. To compare the two measures, we partitioned 80,950 different isoforms of 19,677 genes into deciles by LOEUF score and ran ExtRaINSIGHT separately on the pooled coding sites corresponding to each decile. Again, we computed *ρ* values using INSIGHT together with the *λ_s_* values. We found that both *ρ* and *λ_s_* decreased monotonically with LOEUF decile, with *λ_s_* ranging from 0.28 for the genes having the lowest LOEUF scores to 0.005 for the genes having the highest LOEUF scores, and *ρ* similarly ranging from 0.77 to 0.43 (**Figure 2**). These results suggest that in the 10% of genes under the weakest selection against heterozygous LoF mutations, only 0.5% of sites are subject to ultraselection, but over 40% still experience weaker purifying selection; whereas in the 10% of genes under the strongest selection against LoF mutations, almost 30% of sites are under ultraselection and another ~40% are under weaker purifying selection.

**Figure 2:**
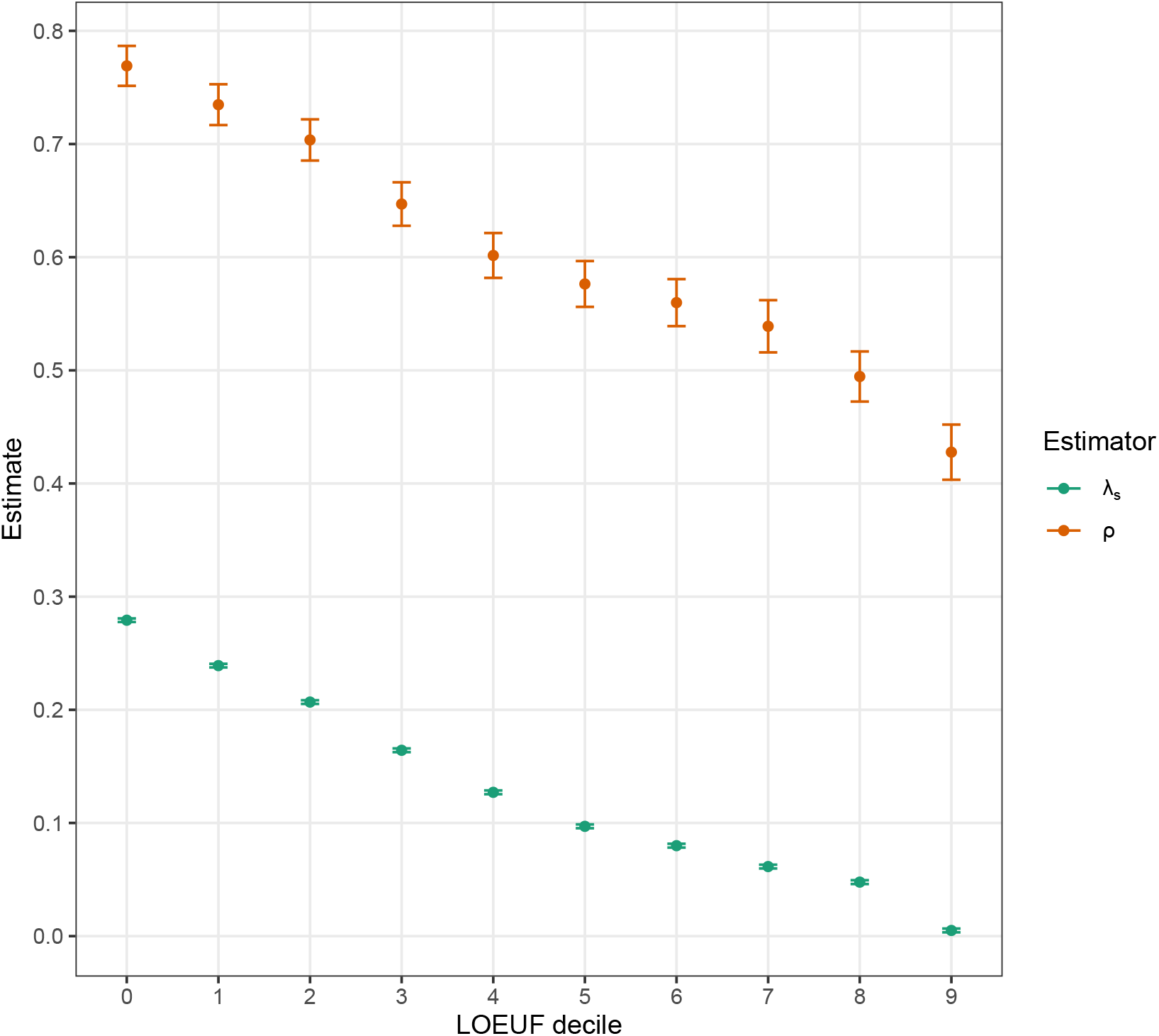
Measures of purifying selection in protein-coding genes by LOEUF decile. The full set of 80,950 isoforms of 19,677 genes was partitioned into deciles according to the loss-of-function observed/expected upper bound fraction (LOEUF) measure [13]. An estimates for each decile is shown for both ExtRaINSIGHT (*λ_s_*) and INSIGHT (*ρ*). Notice that lower LOEUF scores are associated with stronger depletions of LoF variants, so *λ_s_* and *ρ* tend to decrease as LOEUF increases. Error bars indicate one standard error (see **Methods**).

Finally, we considered an alternative grouping of genes by biological pathway, using the top-level annotation from the Reactome pathway database [29] (**Figure 3**). Again, we ran both ExtRaINSIGHT and INSIGHT on each group of genes and observed similar trends in the two measures, with *λ_s_* ranging from 10% to 26%, and *ρ* ranging from 61% to 75%. We found genes annotated as belonging to the “Neuronal System” to be experiencing the most ultraselection (*λ_s_* = 0.26), consistent with other recent findings [9]. Genes annotated as being involved in “Reproduction” showed the least ultraselection (*λ_s_* = 0.10). Notably, the estimates of *λ_s_* exhibited considerably greater variation, as a fraction of the mean, than did estimates of *ρ*. The ratio *λ_s_*/*ρ*—which can be interpreted as the fraction of selected sites experiencing ultraselection— was also highest for “Neuronal System” genes (at 0.36) and lowest for “Reproduction” genes (at 0.17). An analysis of genes exhibiting tissue-specific expression produced similar results, with several brain tissues exhibiting the most ultraselection and vagina exhibiting the least (**Supplemental Fig. S4**).

**Figure 3:**
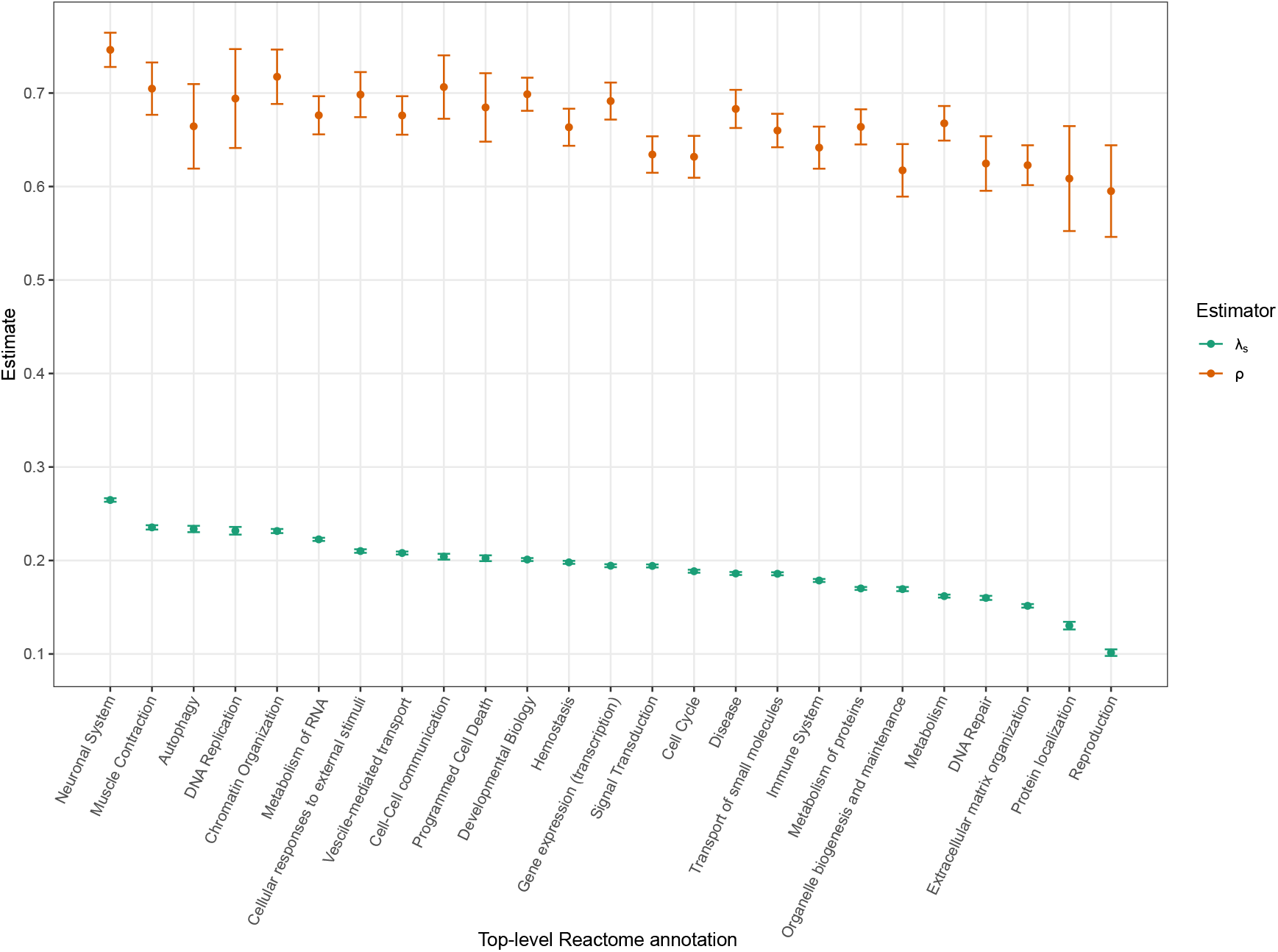
Measures of purifying selection in protein-coding genes by biological pathway. Genes were assigned coarse-grained functional categories using the top-level annotation from the Reactome pathway database [29]. An estimates for each category is shown for both ExtRaINSIGHT (λ_s_) and INSIGHT (*ρ*). Error bars indicate one standard error (see **Methods**).

### Ultraselection in noncoding elements

We carried out a similar analysis on noncoding sequences, including a variety of noncoding RNAs, transcription factor binding sites (TFBS) supported by chromatin-immunoprecipitation-and-sequencing (ChIP-seq) data (from [20]), and unannotated intronic and intergenic regions. Among these sequences, we observed the strongest signature of ultraselection in microRNAs (miRNAs), particularly in evolutionarily “old” miRNAs broadly shared across mammals (designated as “conserved” by TargetScan; see **Methods**), where we estimated *λ_s_* = 0.34 (**Figure 4**). This estimate corresponds to *s*_het_ = 0.016, indicating nearly a 2% reduction in fitness associated with each point mutation in these regions. We found that the seed regions of these miRNAs had even slightly higher values of *λ_s_* = 0.39 (not shown). Interestingly, however, the prevalance of ultraselection was greatly reduced at evolutionarily “new” miRNAs that are not shared across mammals (“nonconserved” in TargetScan), where we estimated only *λ_s_* = 0.031.

**Figure 4:**
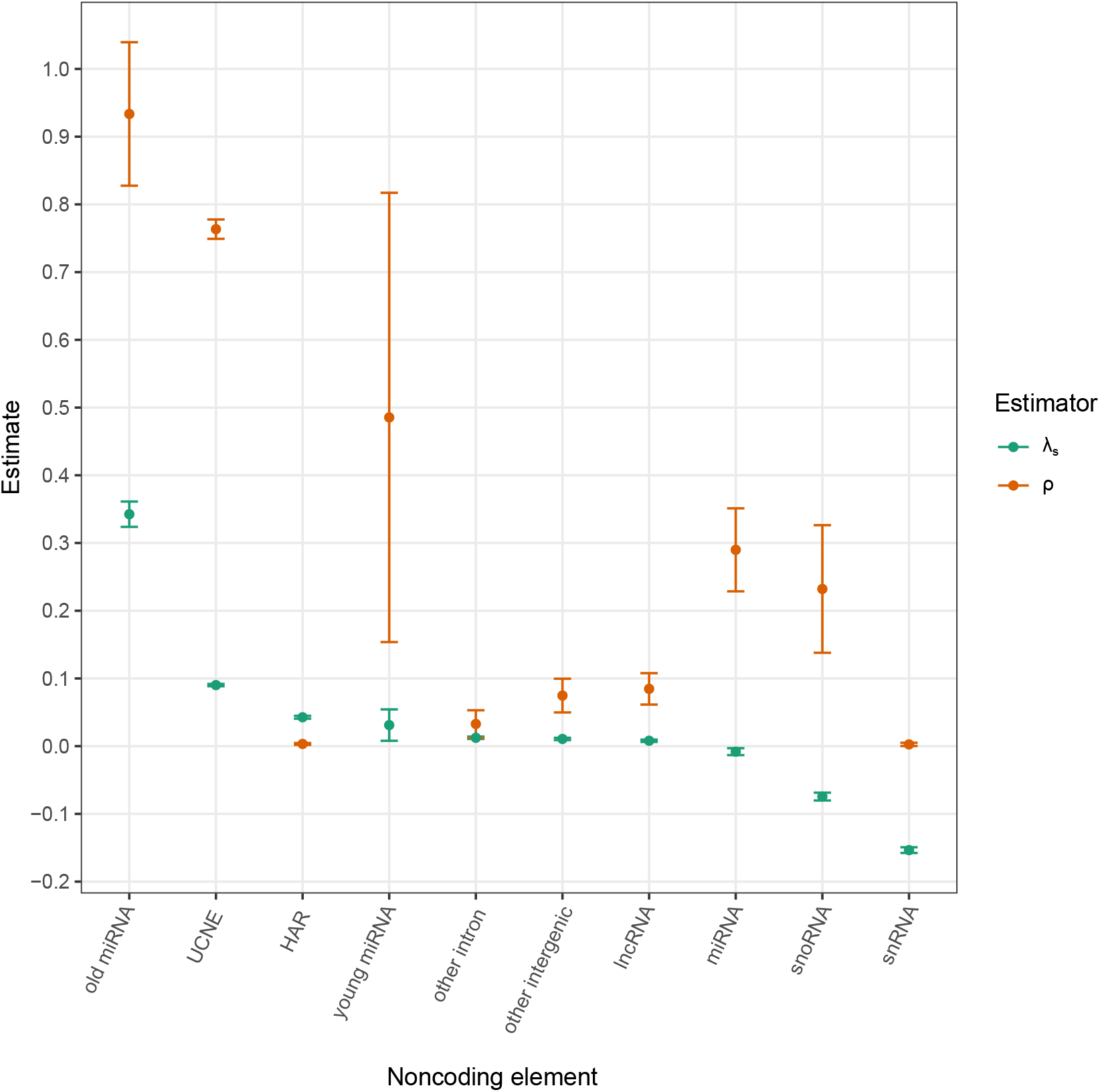
Measures of purifying selection at noncoding elements. Estimates are shown for both ExtRaINSIGHT (*λ_s_*) and INSIGHT (*ρ*). Error bars indicate one standard error (see **Methods**).

Other types of noncoding RNAs also showed little indication of ultraselection: our estimates for long noncoding RNAs (lncRNAs), small nuclear RNAs (snRNAs), and small nucleolar RNAs (snoRNAs) were all close to zero or negative. In an attempt to identify regions within these RNAs that might be subject to stronger selection, we intersected them with conserved elements identified by phastCons [24]. However, we found that even these putatively conserved portions of noncoding RNAs exhibited at most *λ_s_* ≈ 0.05 (in lncRNAs and snRNAs).

When we analyzed a pooled set of all ~2M TFBSs from ref. [20], we obtained a negative estimate of *λ_s_* = –0.08, despite that the same elements yielded a nonnegligible estimate of *ρ* = 0.23. We therefore examined only the binding sites of the 10 TFs whose binding sites showed the largest *ρ* estimates (*ρ* = 0.61 overall; see **Methods**), but even for this putatively conserved set, we obtained an estimate of only *λ_s_* = 0.03. Thus, of the noncoding RNA and TFBSs we considered, only “old” miRNAs appear to experience high levels of ultraselection.

We also evaluated ultraconserved noncoding elements (UCNEs) [30] and noncoding human accelerated regions (HARs) [31–33]—two types of elements that have been widely studied for their unusual patterns of cross-species conservation, and have been shown to function in various ways, including as enhancers [34, 35] and noncoding-RNA transcription units [31]. Interestingly, despite their extreme levels of crossspecies conservation, UCNEs show only modest levels of ultraselection, with *λ_s_* = 0.09. This observation suggests that what is unusual about these elements is not the strength of selection acting on them (which is considerably weaker than that at protein-coding sequences or “old” miRNAs), but instead the uniformity of selection acting at each nucleotide (see **Discussion**). Notably, HARs display only slightly lower levels of ultraselection than UCNEs (*λ_s_* = 0.04) and levels comparable to those of conserved sequences in introns. Thus, despite their rapid evolutionary change during the past 5–7 million years, HARs now appear to contain many nucleotides that are under strong purifying selection in human populations.

### A genome-wide accounting of sites subject to ultraselection

To account genome-wide for the incidence of nearly lethal mutations, we ran ExtRaINSIGHT on a collection of mutually exclusive and exhaustive annotations. For this analysis, we considered CDSs, UTRs, splice sites, lncRNAs, introns, and intergenic regions, but excluded smaller classes of noncoding RNAs, which make negligible genome-wide contributions (**Table 1**). As above, we intersected the lncRNA, intron, and intergenic classes with phastCons elements, and separately considered the conserved and nonconserved partitions of each class. For each category, we multiplied our estimate of *λ_s_* by the number of sites in the category to estimate category-specific expected numbers of sites subject to ultraselection. To account for potential misspecification of the mutational model, we conservatively subtracted from the category-specific estimates of *λ_s_* the estimate for nonconserved intronic regions (0.008). Thus, by construction, the expected number of ultraselected sites in these and similar regions (including nonconserved intergenic and lncRNA sites) was zero.

**Table 1:** Ultraselection across the human genome (based on ExtRaINSIGHT)

| Feature | $\lambda_s$ | ± (stderr)<sup>a</sup> | no. sites (M) | prop. sites | exp. no. (M)<sup>b</sup> | exp. prop.<sup>c</sup> | fold enrich. | exp. lethal<sup>d</sup> | $s_{het}$ |
| --- | --- | --- | --- | --- | --- | --- | --- | --- | --- |
| CDS | 0.149 | 0.002 | 33.8 | 1.18% | 4.8 | 53.1% | 45.2 | 0.11 | - |
| 5' UTR | -0.158 | 0.002 | 8.2 | 0.29% | 0.0 | 0.0% | 0.0 | 0.00 | - |
| 3' UTR | 0.023 | 0.002 | 36.1 | 1.26% | 0.6 | 6.2% | 5.0 | 0.01 | - |
| splice | 0.464 | 0.002 | 0.8 | 0.03% | 0.4 | 4.0% | 146.1 | 0.01 | 2.0% |
| nonconserved lncRNA<sup>e</sup> | 0.008 | 0.002 | 453.6 | 15.78% | 0.0 | 0.0% | 0.0 | 0.00 | - |
| conserved lncRNA<sup>f</sup> | 0.055 | 0.002 | 23.3 | 0.81% | 1.1 | 12.3% | 15.2 | 0.03 | - |
| nonconserved intron<sup>e</sup> | 0.008 | 0.002 | 972.6 | 33.83% | 0.0 | 0.0% | 0.0 | 0.0 | - |
| conserved intron<sup>f</sup> | 0.057 | 0.002 | 44.3 | 1.54% | 2.2 | 24.4% | 15.8 | 0.05 | - |
| nonconserved intergenic<sup>e</sup> | 0.003 | 0.002 | 1255.5 | 43.67% | 0.0 | 0.0% | 0.0 | 0.00 | - |
| conserved intergenic<sup>f</sup> | 0.051 | 0.002 | 46.9 | 1.63% | 2.0 | 22.4% | 13.7 | 0.05 | - |
| Total |  |  | 2875.1 | 100.00% | 9.0 | 100.0% |  | 0.26 |  |

<sup>a</sup>The similar values of the standard errors (equal after rounding) reflect the maximum of 1M sites used for estimation.
<sup>b</sup>Expected number of ultraselected sites after adjusting for background. In this case, the estimate for nonconserved introns (0.008) was subtracted from each estimate of $\lambda_s$ (see **Supplemental Table S1** for a less conservative correction).
<sup>c</sup>Expected proportion of ultraselected sites after adjusting for background.
<sup>d</sup>Expected number of new lethal or nearly lethal mutations per diploid individual, assuming a mutation rate of $1.2 \times 10^{-8}$ per generation per site.
<sup>e</sup>Sites not classified as conserved by phastCons.
<sup>f</sup>Sites classified as conserved by phastCons.

Overall, we estimated that 0.31% of the human genome is ultraselected, with 53% of ultraselected sites falling in CDSs, 24% in conserved introns, 22% in conserved intergenic regions, 12% in conserved lncRNAs, 6% in 3’ UTRs and 4% in splice sites. Notably, ultraselected sites are overrepresented 45-fold in CDSs, but CDSs still account for only about half of ultraselected sites. Splice sites are overrepresented 146-fold but make a minor overall contribution owing to their small number.

Our assumption is that any point mutation at these ultraselected sites will be nearly lethal, and simulations indicate that the detected sites are indeed subject to extreme purifying selection (see **Discussion**). Thus, if we multiply the expected numbers of sites by twice (allowing for heterozygous mutations) the estimated per-generation, per-nucleotide mutation rate (here assumed to be 1.2 × 10^-8^ [36]), we obtain expected numbers of *de novo* nearly lethal mutations per potential zygote (“potential” because some mutations will act prior to fertilization). By this method, we estimate 0.26 nearly lethal mutations per potential zygote. By construction, these nearly lethal mutations occur in the same category-specific proportions as the ultraselected sites (53% from CDS, 24% from introns, etc.). Thus, we expect 0.11 nearly lethal coding mutations per potential zygote and another 0.15 such mutations at various noncoding sites.

If we carry out a less conservative version of these calculations, by subtracting the *λ_s_* estimate for nonconserved intergenic regions (0.003) rather than the one for intronic regions, we estimate 0.54% of the genome to be ultraselected, with 32% falling in CDSs (**Supplemental Table S1**). The expected number of nearly lethal mutations per potential zygote increases to 0.43, of which 0.12 fall in CDSs. Taking these calculations together, we estimate a range of 0.26–0.43 nearly lethal mutations per potential zygote, implying a high genetic burden but one that appears to be roughly compatible with other lines of evidence (see **Discussion**).

We performed a parallel analysis using INSIGHT, to estimate the numbers and distribution of more weakly deleterious mutations (**Table 2**). In this case, we estimate that 3.2% of sites are under selection and the expected number of *de novo* deleterious mutations per fertilization is 2.21. The fraction of deleterious mutations from CDS is 22%, with most of the remainder coming from introns and intergenic regions. lncRNAs and 3’ UTRs also make significant contributions. Taking the ExtRaINSIGHT and INSIGHT estimates together, we estimate that each potential fertilization event is associated with 0.26–0.43 new lethal mutations and an additional 1.78–1.94 new mutations that are more weakly deleterious. One way to interpret these numbers is that, conditional on a threshold level of fitness (i.e., the existence of no nearly lethal mutations), each person contains an expected ~2 new mutations that are sufficiently strongly deleterious that they would tend to be eliminated from the population on the time-scale of human-chimpanzee divergence (as measured by INSIGHT), at least if humans continued to experience historical levels of purifying selection. That person’s genetic load would derive from both these new mutations and similar weakly deleterious mutations passed down from his or her ancestors.

**Table 2:**
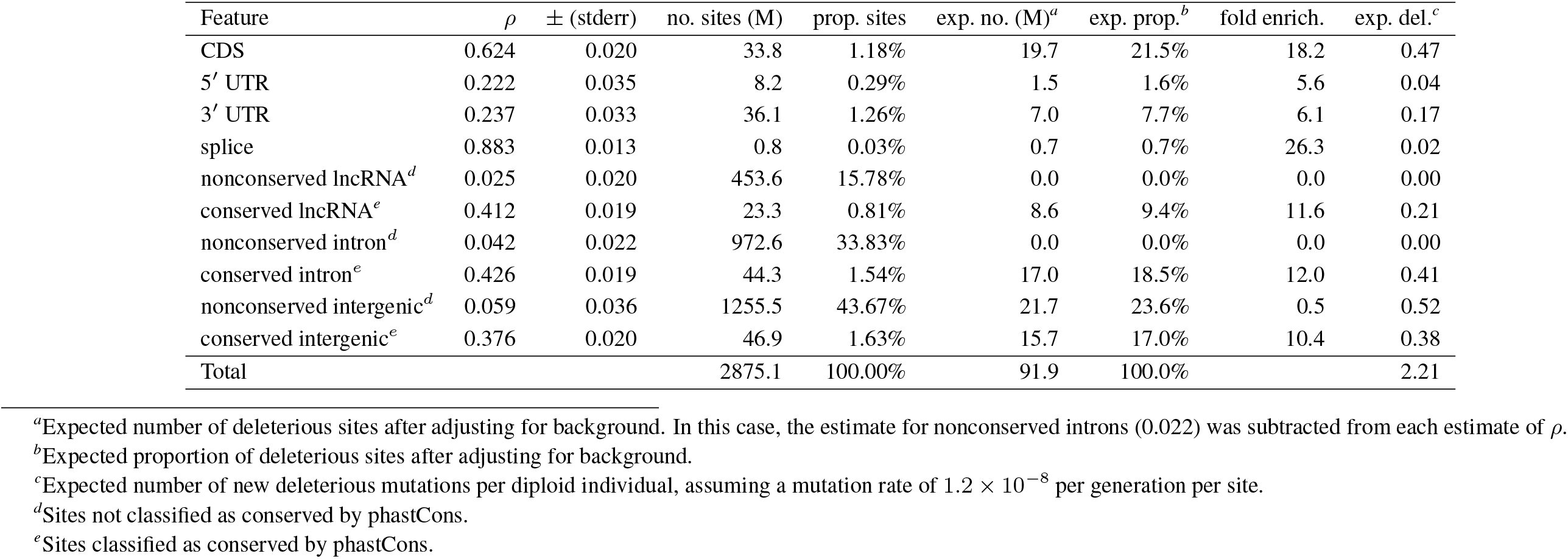
Weaker selection across the human genome (based on INSIGHT)

### Persistent misspecification of the mutation model at promoter regions and TFBSs

As noted above, we observed a consistent tendency to estimate negative values of *λ_s_* at the 5’ ends of genes, including in 5’ UTRs and core promoters (**Figure 1**), as well as at TFBSs and some noncoding RNAs from across the genome (**Figure 4**). In an attempt to bound the genomic regions near protein-coding genes that give rise to these negative estimates, we applied ExtRaINSIGHT in a series of windows near the 5’ and 3’ ends of genes, pooling data from all ~20,000 genes (**Figure 5a**). We found that the effect was most pronounced in the 5’ UTR, where we estimated *λ_s_* = –0.16 (see **Figure 1**) and in the 250bp immediately upstream of the TSS (*λ_s_* = –0.13). As we looked farther upstream, it diminished fairly rapidly, with *λ_s_* = –0.05 in the (–500, –250) window and *λ_s_* = –0.02 in the (–1000, –500) window. By the (–2000, –1000) window, the estimates had returned to slightly positive values. We did not observe negative estimates near the 3’ ends of genes, and the estimate for 4d sites within the CDS was only slightly negative. Therefore, the tendency to estimate *λ_s_* < 0 near genes appears to be limited to the 5’ UTR and the ~1kb region upstream of the TSS.

**Figure 5:**
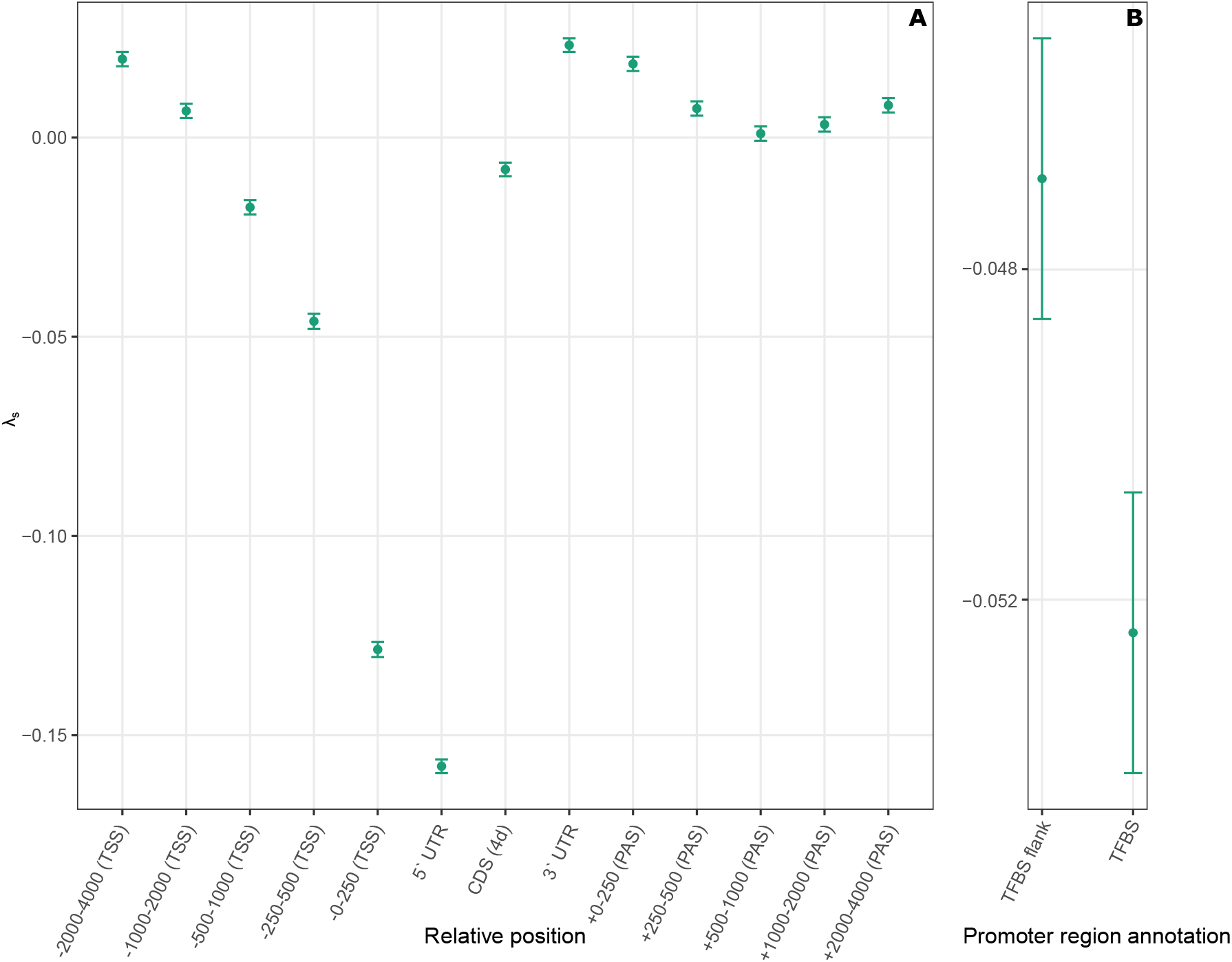
Ultraselection in genomic intervals upstream and downstream of protein-coding genes. (**A**) Windows upstream of the transcription start site (TSS) and downstream of the polyadenylation site (PAS) are labeled on the *x*-axis. The 5′ and 3′ UTRs are also shown, as are fourfold degenerate (4d) coding sites (CDS). Estimates of *λ_s_* with error bars indicating one standard error are shown on the *y*-axis. (**B**) Estimates for the extended promoter region (2kb upstream of the TSS) within transcription factor binding sites (TFBS) annotated in the Ensembl Regulatory Build [42] and in the immediate flanking sequences (10bp on each side). The difference is highly statistically significant by a likelihood ratio test based on the ExtRaINSIGHT likelihood model (*p* = 2.8 × 10^-13^).

We hypothesized that, despite being well-calibrated across the majority of the genome (**Supplemental Fig. S1**), our mutation model is misspecified in promoter regions, perhaps owing to correlations of mutation rates with features such as chromatin accessibility or hypomethylation. We therefore adapted our model to consider the predicted state from an application of the 25-state ChromHMM model [37, 38] to Roadmap Epigenomics data [39] as a categorical covariate and refitted it to the data, trying ChromHMM predictions for several cell types. However, we found that this approach did not eliminate the tendency for negative estimates of *λ_s_* (results not shown), perhaps because the available epigenomic data has too coarse a resolution or is not well matched by cell type.

Having observed negative estimates of *λ_s_* also at TFBSs outside of promoter regions, however, we wondered if the effect could be driven, at least in part, by TF binding itself, which has been shown to be mutagenic in melanoma [40, 41]. In an attempt to isolate the effects of TF binding, we applied ExtRaINSIGHT separately to predicted TFBS in extended promoter regions, using predictions from the Ensembl Regulatory Build [42], and to the immediate flanking 10bp on either side of these predictions, excluding flanking sequences that themselves included TFBSs. Interestingly, we found that estimates of *λ_s_* were significantly more negative in the TFBSs than in the immediate flanking sites (**Figure 5b**; *p* = 2.8 × 10^-13^, likelihood ratio test), suggesting a possible influence from the mutagenic effects of TF binding (see **Discussion**). In the end, we were not able to eliminate this apparent problem with our mutation model, but its effects appear to be generally quite local to TSSs and TFBSs and therefore are likely to have a limited impact on our genome-wide analyses.

## Discussion

In this article, we have introduced a new method, called ExtRaINSIGHT, for measuring the prevalence of strong purifying selection, or “ultraselection,” on any collection of sites in the human genome, including noncoding as well as coding sites. ExtRaINSIGHT enables maximum-likelihood estimation of a parameter, denoted *λ_s_*, that represents the fractional depletion in rare variants in a target set of sites relative to matched “neutral” sites, after accounting for neighbor-dependence and local variation in mutation rate. We have shown that when *λ_s_* is sufficiently large (approximately >0.2 for our data) and mutation-selection balance is assumed, 1 – *λ_s_* is expected to have an inverse relationship with the selection coefficient against heterozygous mutations, 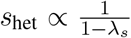, which allows *s*_het_ to be estimated for a target collection of sites. Simulations indicate that this approximation is reasonably good, although it is biased downward when selection is variable across sites (see **Methods, Supplemental Fig. S3**) and biased upward near the boundary of *λ_s_* ≈ 0.2 (**Supplemental Fig. S2**). We have surveyed the prevalence of ultraselection in both coding and non-coding regions of the human genome and found it to be particularly strong in splice sites, 0-fold degenerate (0d) coding sites, and evolutionarily ancient miRNAs. On the other hand, ultraselection is mostly absent in other noncoding RNAs, untranslated regions of protein-coding genes, and transcription factor binding sites, as well as in fourfold degenerate (4d) coding sites. We have also shown that neural-related genes and genes expressed in the brain are enriched for large estimates of *λ_s_* in their codings sequences, whereas reproduction-related genes are enriched for small estimates of *λ_s_*.

Interestingly, we found only a modest prevalence of ultraselection in ultraconserved noncoding elements (UCNEs), despite their near-complete sequence conservation over hundreds of millions of years of evolution [30]. It has been suggested that this extreme conservation is indicative of strong purifying selection (e.g., [30]), although most such observations have not been accompanied by direct estimation of selection coefficients. One exception is an early study by Katzman et al. [43], where ultraconserved elements in humans were estimated to be experiencing substantially stronger selection (by about 3-fold) than nonsynonymous sites in protein-coding sequences, although the absolute strength of selection was estimated to be modest (mean of 2*N_e_s* ≈ –5) and the analysis was based on only 72 individuals. The assumption of strong levels of selection has been difficult to reconcile with observations that organisms often appear to function normally after deletion of UCNEs, as when complete deletion in mice of megabase-long gene deserts containing UCNEs failed to produce detectable phenotypes [44]. More recently, Snetkova et al. found that UCNEs were remarkably resilient to mutation, with a majority continuing to function as enhancers in transgenic mouse reporter assays even after being subjected to substantial levels of mutagenesis [45]. Our observations suggest that these apparently contradictory observations—high sequence conservation and resilience to mutation—can be reconciled if UCNEs are predominantly under relatively weak selection, that is, selection strong enough to prohibit fixation of new mutations on the time scales of interspecies divergence but weak enough that rare variants are not substantially depleted. Indeed, we find considerably lower levels of ultraselection in UCNEs (*λ_s_* = 0.09) than in 0d sites in coding regions (*λ_s_* = 0.22) or in ancient miRNAs (*λ_s_* = 0.34). At the same time, these classes of sites tend not to show perfect conservation in cross-species comparisons, primarily because they tend to be interspersed with less conserved sites (e.g., 4d sites or non-pairing sites in miRNAs). Thus, what seems to be most unusual about UCNEs is not the extreme level of purifying selection they experience but rather the uniformity of purifying selection across hundreds of bases. In most cases it is still unknown what causes this uniformity, although it has been speculated that it may result from overlapping functional roles, such as overlapping binding sites, structural RNAs, and coding regions [30].

It is instructive to compare our estimates of *s*_het_ with Cassa et al.’s [11] mean estimate of *s*_het_ = 0.059 for predicted loss-of-function (pLoF) variants in protein-coding genes. Our estimate for splice sites (*λ_s_* = 0.46, *s*_het_ = 0.020) is reasonably concordant with this estimate, assuming that many but not all splice-site-disrupting mutations result in loss of function, and allowing for our possible underestimation of *s*_het_ in the presence of variability across sites. Our estimate of *λ_s_* = 0.22, *s*_het_ = 0.014 for missense mutations at 0d sites is plausible—e.g., it is roughly comparable with experimentally derived estimates for *s*_het_ of 1–3% for strongly deleterious mutations in yeast and flies [11, 46, 47]—but it seems at first glance to be high in comparison to Cassa et al’s pLoF estimates, given that a majority of missense mutations are presumably neutral or only mildly deleterious.

Studies based on the site-frequency-spectrum have tended to infer long-tailed distributions—such as gamma or lognormal distributions—for the DFE for new amino-acid replacements, often augmented with point-masses at zero [5–8]. The best-fitting such model in a representative recent study by Kim et al. [8], based on a fairly large sample size (432 Europeans from the 1000 Genomes Project), implied a mean selection coefficient against amino-acid replacements of s = 0.007. These methods assume additivity, so this estimate corresponds to only 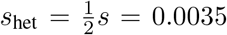, about one fourth of our estimate for 0d sites. It is therefore possible that our estimate is too large, particularly since it falls near the boundary of the regime where mutation-selection balance holds. At the same time, it is also possible that these SFS-based methods have systematically underestimated the weight of the tail of the DFE, which is well known to be difficult to measure based on the SFS and samples of modest size (e.g., [7]). Notably, if we apply ExtRaINSIGHT to data simulated under Kim et al.’s DFE, we obtain an estimate of only *λ_s_* = 0.04, compared with *λ_s_* = 0.22 for real 0d sites (**Supplemental Table S2, Supplemental Fig. S5).** Thus, the patterns of rare variants present in the deeply sequenced gnomAD data set do not seem to be consistent with the DFEs inferred from smaller data sets, likely because these inferred DFEs have failed to accurately describe the tail of the distribution. It therefore seems plausible that our fourfold higher estimate of *s*_het_ ≈ 1.4% is closer to the true mean value than these SFS-based estimates.

One particular challenge with our method is accommodating variation across sites in shet. Because our likelihood function is based simply on the presence or absence of rare variants across a collection of exhangeable sites, it carries limited information about the second moment of the DFE. Unlike ref. [11], we cannot aggregate together all mutations likely to result in loss-of-function of a gene, which permits inference of the genewise distribution of *s*_het_. Notably, however (see **Methods**), in the presence of variation in *s*_het_, our approximate estimator will describe the harmonic mean, rather than the arithmetic mean, of the true values. Consequently, it will have a predictable downward bias, meaning that it can be interpreted as a lower-bound on the true arithmetic mean. This downward bias is consistent with our observations in splice sites. For 0d sites and ancient miRNAs, it provides additional confidence in our seemingly high estimates, suggesting that the true values could be even larger. It may be possible in future work to extend our methods to consider a distribution of shet values, for example, by introducing a scheme for grouping sites into elements analogous to the genes in ref. [11].

Another possible concern with our approach is that, in estimating *λ_s_* from the rare variants missing from the target sites, ExtRaINSIGHT inevitably will pick up not only on strong selection against nearly lethal mutations but also, to a degree, on selection on a large class of more weakly deleterious mutations. Even if these more weakly deleterious mutations are inefficiently eliminated over the short time scale relevant for rare variants, their cumulative effect could still be substantial relative to that from strongly deleterious mutations if they are much larger in number—which is plausible if the weight in the tail of the true DFE is not too large. Such a scenario could potentially lead to overestimation of *λ_s_* and, consequently, of *s*_het_ and of the numbers of nearly lethal mutations per potential fertilization.

We attempted to examine this question by simulating data under four different DFEs, representing scenarios from quite weak selection (as we observe in TFBSs) to quite strong selection (as we observe at evolutionarily ancient miRNAs), applying ExtRaINSIGHT to the simulated data, and then decomposing the DFE into a component associated with the rare variants removed by selection and a component associated with the remaining rare variants (which we can trace in simulation; see **Supplemental Fig. S5** and **Supplemental Table S2**). We found, overall, that the missing variants detected by ExtRaINSIGHT are strongly enriched for strong purifying selection. In the case of quite strong selection (similar to what we infer at 0d sites or miRNAs), they predominantly have *s*_het_ > 0.01, with mean values of *s*_het_ ≈ 0.03. Even in the case of Kim et al.’s inferred DFE (which, as discussed above, may underestimate the tail), the mean *s*_het_ = 0.03 for the missing rare variants, although in this case substantially more of them have *s*_het_ < 0.01. Overall, we find that, with mean *s*_het_ ≈ 0.03, these rare variants are indeed under quite strong purifying selection, although our power to separate strong selection from nearly neutral evolution does depend on the original DFE. At this selection coefficient, some rare variants may persist for a few generations, but, according to Kimura and Ohta’s [48] formulas, the expected number of generations until extinction will be no more than about half of the neutral expectation, which itself is quite low (see **Supplemental Text**). Thus, it seems reasonable to regard these variants as “nearly lethal.”

What are the implications of our estimate of ~0.3–0.4 for the number of nearly lethal mutations per potential fertilization? This estimate implies a fairly high genetic burden for severely deleterious mutations (not to mention the additional burden imposed by weakly deleterious mutations), but one that appears to be in the plausible range (e.g., [23, 28]). One rough point of comparison is the rate of spontaneous abortion, which has been estimated to be as high as 50% for mothers of prime reproductive age [49, 50]. This quantity, of course, is not the same as the rate of nearly lethal mutations, for a variety of reasons—spontaneous abortion typically describes death prior to birth conditional on a detectable pregnancy, whereas our measure includes mutations that are lethal near the time of fertilization or even prior to fertilization, and also includes mutations that cause death after birth, that do not cause death but prevent an organism from reproducing, or that severely reduce fitness over several generations. In addition, many of the mutations that cause spontaneous abortion in the fetus are not point mutations, but instead major structural variants that often alter karyotype [49]. At the same time, spontaneous abortion is only partly a consequence of the genetics of the embryo, also depending strongly on the environment and the genetics of the mother. Nevertheless, it is notable that these quite different estimates are in rough agreement with one another, suggesting an overlap in what they are measuring, perhaps with other factors approximately cancelling.

Throughout this article, we have compared *λ_s_* estimates from ExtRaINSIGHT with *ρ* estimates from INSIGHT, in order to evaluate the relative fractions of sites subject to ultraselection and weaker forms of purifying selection. It is worth noting, however, that the two methods are not based on precisely the same assumptions and therefore are not exactly comparable. Unlike ExtRaINSIGHT, INSIGHT measures natural selection on the time scale of the human-chimpanzee divergence (5–7 MY), assuming that functional roles are relatively constant during that time period. It also incorporates positive selection as well as purifying selection into its model, although positive selection appears to make at most a minor contribution to *ρ* in this setting (see **Methods**). Finally, INSIGHT makes use of a much simpler Jukes-Cantor mutation model, with no accounting for neighbor-dependence in mutation rate (although it does account for regional variation across the genome). As a result, differences between *λ_s_* and *ρ* could result in part from matters such as gain and loss of functional elements on human/chimp time scales, misspecification of the Jukes-Cantor mutation model, or contributions from positive selection. Nevertheless, we expect these differences to have relatively minor effects, and the estimates from INSIGHT and ExtRaINSIGHT appear to be fairly consistent overall, with *ρ* and *λ_s_* well correlated but *ρ* > *λ_s_* in all cases. Therefore, we believe it is reasonable to approximately characterize the DFE by treating *λ_s_* as a measure of ultraselection and the difference *λ_s_* – *ρ* as a measure of selection that is weaker but sufficiently strong to result in removal of deleterious variants on the time scale of human/chimpanzee divergence.

While our mutation model fits the data well across most of the genome, we were not able to eliminate an apparent misspecification of this model in promoter regions as well as at other TFBSs and at some noncoding RNAs. This misspecification is unlikely to be explained by unusual base or word composition in these regions, nor by regional variation in overall mutation rate, because these features are explicitly addressed by our model. We also could not eliminate it by explicitly conditioning on chromatin state, using the ChromHMM model [37, 38], although it is possible that our approach was limited by the resolution and cell-type-specificity of the available epigenomic data. Interestingly, the best predictor we could identify for elevated mutation rates was TF binding itself. There is accumulating evidence from melanoma that TF binding may be mutagenic, likely because it interferes with DNA repair [40, 41], so it seems possible that TF binding is, at least in part, a driver of elevated germ-line mutation rates in these regions. It is worth noting that if TF binding indeed itself significantly alters mutation rates, this phenomenon would considerably complicate efforts to measure natural selection on TFBS, which is generally accomplished by contrasting rates of polymorphism and/or divergence within binding sites relative to nearby flanking sites, under the assumption that mutation rates are approximately equal in these regions (e.g., [20, 26, 51]). However, the strength of this mutagenic effect in the germline remains unknown, and unless it is particularly pronounced, it likely has a minor effect on analyses at longer evolutionary time scales, where natural selection probably dominates in determining patterns of polymorphism and divergence. In any case, more work will be needed to develop a full understanding of these potential mutational biases and account for them in analyses of selection on binding sites.

## Methods

### Data for neutral model

The data for our neutral model consisted of rare variants (MAF <0.001) from gnomAD (v3) within the genomic regions identified by Arbiza et al. [20] as putatively free from selection, unduplicated, non-repetitive, and reliably mappable. These regions were mapped to the hg38 human assembly using liftOver [52]. We further removed all CpG sites, which we expected to be difficult to model owing to methylation-induced hypermutation, and all sites having an an average sequencing coverage across individuals of < 20 reads.

### Mutation model

To fit the mutation model to these putatively neutral sites, we first calculated the relative frequencies of each type of mutation *a* → *b* and of the absence of a mutation (*a* → *a*), conditional on the identities of a, b, and the three flanking nucleotides on each side. This required collecting 4^8^ = 65536 distinct counts (minus the excluded CpGs) and normalizing them to sum to one separately for each a and flanking nucleotides. We then obtained adjusted rates by combining the (logits of) these raw relative rates with a collection of covariates likely to be correlated with real or apparent rates of mutation in a linear-logistic model. In particular, we used four covariates: the raw relative frequency, the logarithm of the reported average sequencing coverage from gnomAD, the fractional G+C content in a 200bp window, and an indicator for whether or not each site fell in a CpG island (based on the UCSC Genome Browser track of the same name [52]). We fitted this model to the observed rates of mutation at variable and nonvariable sites, sampling 1% of putatively neutral sites for efficiency. Finally, we further adjusted the estimated rates for regional variation in mutation rate by sliding a 150kb window along the genome in 50kb increments, and fitting a linear-logistic model to the neutral sites in each window, with the logit of the previously estimated rate as a covariate with coefficient one and a free intercept term, which could be interpreted as a local scaling factor. Together, these steps allowed us to estimate an absolute rate for the emergence of each allele at each site in the genome. When we compare the predicted rates with actual rates within the neutral regions, we can see that the model is quite well calibrated (Supplemental Fig. S1).

### Approximate model for ultraselection

Following equation 1, the log likelihood function is given by,

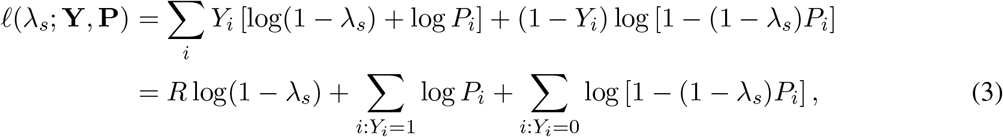

where 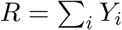 is the number of rare variants. When the *P_i_* values are small (as is typical), it is possible to obtain a reasonably good closed-form estimator for *λ_s_* by making use of the approximation log(1–*x*) ≈ –*x*. In this case,

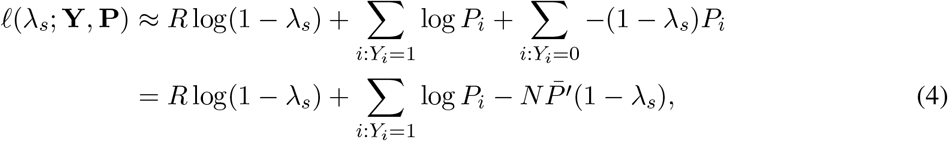

where 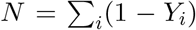 is the number of invariant sites and 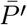 is the average value of *P_i_* at the invariant sites. It is easy to show that this approximate log likelihood is maximized at,

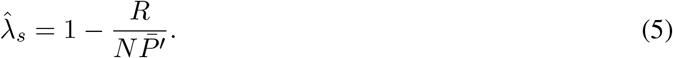

However, this procedure leads to a biased estimator for *λ_s_*. A correction for the bias leads to the following, intuitively simple, unbiased estimator:

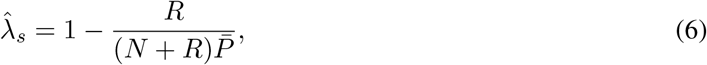

where 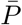 is the average value of *P_i_* at all sites. In other words, 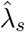 is given by 1 minus the observed number of rare variants divided by the expected number of rare variants under neutrality, which is simply the total number of sites, *N* + *R*, multiplied by the average rate at which rare variants appear, 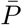.

### Full allele-specific model

In practice, we use a model that distinguishes among the alternative alleles at each site and exploits our allelespecific mutation rates. This model behaves similarly to the simpler one described above, but yields slightly more precise estimates, because the mutation rates for different alternative alleles can differ appreciably, and because multiple alternative alleles are often present at a single site in the gnomAD data.

In the full model, we assume separate indicator variables, 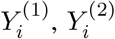, and 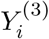, for the three possible allele-specific rare variants at each site, and corresponding allele-specific rates of occurrence, 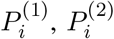, and 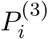. We further make the assumption that the different rare variants appear independently. Thus, the likelihood function generalizes to (cf. equation 1),

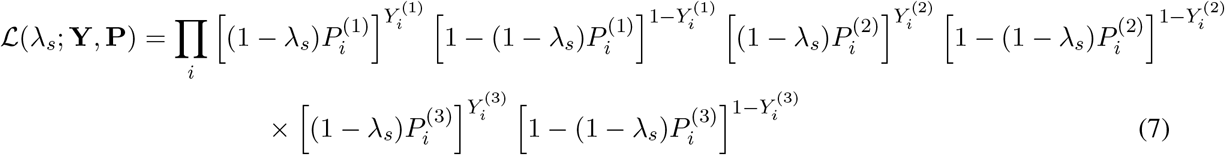

where we redefine 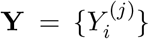 and 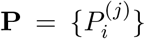 for *j* ∈ {1, 2, 3}. Notice that, when more than one alternative allele is present, 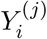 will be 1 for more than one value of *j*.

ExtRaINSIGHT simply maximizes this function with respect to *λ_s_* numerically. To improve efficiency, it considers at most one million sites, subsampling down to one million if more are provided. Standard errors for *λ_s_* are estimated by taking the square root of the inverse of the negative second derivative of the log likelihood function. ExtRaINSIGHT also reports a p-value based on a likelihood ratio test of an alternative hypothesis of *λ_s_* = 0 relative to a null hypothesis of *λ_s_* = 0, assuming twice the log likelihood ratio has an asymptotic *χ*^2^ distribution with one degree of freedom under the null hypothesis.

### Relationship between *s*_het_ and *λ*_s_ under mutation-selection balance

When selection against heterozygotes is strong, the equilibrium allele frequency at mutation-selection balance is given by 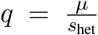 (reviewed in [17]). The frequency of mutant alleles in a random sample of 2*N* chromosomes (where *N* is the number of diploid individuals) will be Poisson-distributed with mean 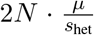 (c.f. [11]), and the expected number of polymorphic sites in a collection of *M* sites is *E*[*X*] = *M*(1 – *e*^-2*Nμ*/*s*_het_^). Ignoring common variants for the moment, the same expectation under the ExtRaINSIGHT model is given by 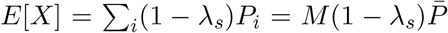, where *P* is the mean value of *P_i_* over the sites in question. By setting these quantities equal to one another, we obtain,

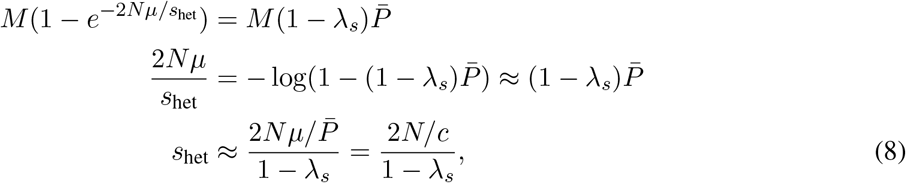

where 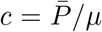. With our data, we find that 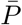 varies little from one set of sites to another, hovering close to 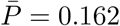. Assuming *μ* = 1.2 × 10^-8^, we obtain *c* = 1.35 × 10^7^.

This derivation can be adjusted to accommodate common variants (with MAF > 0.001, under our assumptions), but this correction has little effect in practice with our data, because only about 3% of variants are common. Since the relationship is approximate anyway, we use the simpler version above.

It is instructive also to consider the case where *s*_het_ varies across sites. In this case, if *s_i_* is the selection coefficient against heterozygotes at site *i* and if each *s_i_* is sufficiently strong for mutation-selection balance to hold, then,

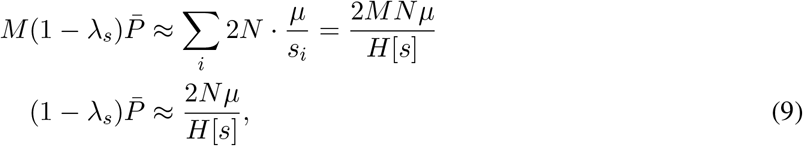

where 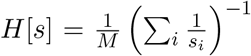 is the harmonic mean of the *s_i_* values. This relationship is equivalent to the one above but with *H*[*s*] in place of *s*_het_. Therefore, in this case, equation 8 yields an estimator not for the arithmetic mean, but for the harmonic mean of the variable s_i_ values across sites. It will therefore tend to underestimate the arithmetic mean in the presence of variable selection. This observation provides an explanation for the downward bias observed in **Supplemental Fig. S1**.

A further generalization of interest is to assume that a fraction *π*_0_ of the sites of interest are not under selection at all. In this case, the rare variants will arise as a mixture of sites under selection (and at mutationselection balance) and sites at which the neutral rate applies. Thus,

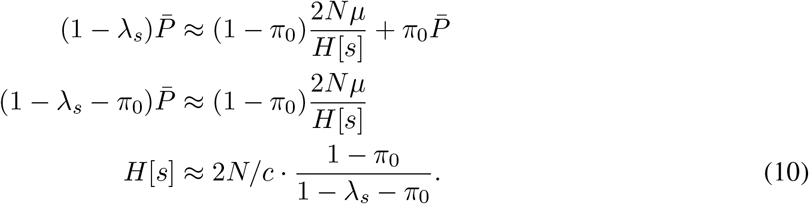

Consequently, if the sites of interest are known to include a component of neutrally evolving sites, and if the 5 fraction *π*_0_ can be estimated, then a portion of the downward bias in estimation of the selection coefficient can be removed. In particular, the quantity *ρ* estimated by INSIGHT should function as a fairly good estimate of 1 – *π*_0_. Therefore, if estimates of 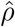 and 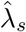 are both available, one can obtain an adjusted estimate of the harmonic mean of *s* as,

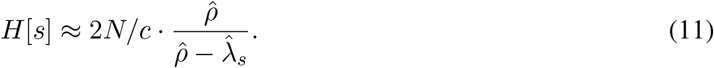

### Calculating the fraction of sites under selection using INSIGHT

To estimate the total fraction of sites under selection we applied INSIGHT [19, 20] in parallel to ExtRaINSIGHT, using the same sets of foreground and background (“neutral”) sites. INSIGHT reports a maximum-likelihood estimate of a quantity *ρ* that measures the fraction of all sites subject to selection on the time scale of the human-chimpanzee divergence (5–7 MY). This quantity includes sites under positive selection as well as those under purifying selection, but for large collections of sites in the human genome the contribution of positive selection is generally negligible (see [20, 53]). For efficiency, we used a re-engineered version of INSIGHT, called INSIGHT2, that is mathematically equivalent to the original but performs numerical optimization using the BFGS algorithm rather than expectation maximization [54]. INSIGHT2 is currently only available for the hg19 assembly so we first mapped annotations from hg38 to hg19 using liftOver, ignoring sites outside of regions of one-to-one mapping. As with ExtRaINSIGHT, we randomly sampled one million sites from larger data sets, to improve efficiency. Notably, INSIGHT makes use of data from Complete Genomics rather than the gnomAD data set for allele-frequency information (see [20]). INSIGHT calculates the standard error of its estimates of *ρ* by taking the inverse of the corresponding diagonal term of the negative Hessian matrix of the log likelihood function at the MLE.

### Genomic annotations and data processing

Annotations for CDS, 5′ UTR, 3′ UTR, and introns were defined using the ensembldb Bioconductor pack-age, which interfaces directly with Ensembl. We included only autosomal protein-coding genes. Splice sites were defined as the two nucleotide sites at each of the 5′ and 3′ ends of introns. Within the promotor regions, we used the Ensembl Regulatory Build to locate transcription factor binding sites, which are inferred from experimental data. Flanking regions of TFBS were defined as the 10 bases on either side of each TFBS. We obtained annotations for lncRNA, snRNA, snoRNA, miRNA also using ensembldb, again restricting them to the autosomes. For all of these annotations, we excluded any regions included in the CDS annotations.

Human accelerated regions (HARs) were obtained from Supplemental Table 1 of ref. [55], a compilation from five previous studies. Ultraconserved noncoding elements (UCNEs) were obtained from UCNEbase [56]. These HARs and UCNEs were defined with respect to hg19, so we mapped them to hg38 using liftOver.

Functional categories were obtained from the Reactome database [29], considering only “top-level” human terms that included at least 100 genes. Tissue specific genes expression data were obtained from Supplemental Table 1 in ref. [57]. Genes were classified as tissue-specific if they had a TS score of greater than three, indicating that they are expressed in that tissue at a level roughly 2^3^ times as high as the average expression level in all other tissues. Note that this definition allows a gene to be “tissue-specific” in more than one tissue. For each category of interest (based on pathway or gene expression), we applied ExtRaINSIGHT to the union of CDS exons of all associated protein-coding gene.

### Simulations

To test our ability to estimate *s*_het_ from *λ_s_* (as shown in **Supplemental Fig. S3**), we conducted simulations under a realistic demographic model and various “true” values of shet. We then estimated *λ_s_* for each data set, converted *λ_s_* to shet via equation 2, and compared this estimate to the true value. In each case, we used the simulator developed by Weghorn et al. [18] to generate 100,000 independent nucleotide sites for a population of 71,702 diploid individuals with bottlenecks and growth patterns matching based on a European demographic history. We carried out an initial round of simulations assuming a constant value of *s*_het_ per simulated data set, with *s*_het_ ranging from 0.0001 to 0.5, and a second round in which sitewise values of shet were drawn from an exponential distribution with a mean equal to each of the same values. When applying equation 2, we used the mean rate of rare variant occurrence, 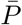, observed in each simulated data set, which tended to be similar, but not identical, to that from the real data. Similarly we used the mutation rate employed in the simulations (2.2 × 10^-8^ per generation per site), which had been adjusted upward to make the frequency of rare variants in the simulated data similar to that in the real data.

In a second series of experiments, we simulated data from DFEs based on real data and evaluated the DFE associated with the “missing” rare variants measured by ExtRaINSIGHT, as well as the quality of the *λ_s_* and *s*_het_ estimators (**Supplemental Table S2** and **Supplemental Fig. S3**). We used four DFEs: (1) one derived from ref. [8] based on data from the 1000 Genomes Project, consisting of a mixture of a point-mass at zero (3.1% weight) and a Gamma distribution with α=0.1930 and θ=0.0168 (“Kim et al.” in **Table S2**); (2) a version of the same DFE with a larger value of the shape parameter (*α* = 0.87) to better mimic the patterns we observed at 0d sites (“0d CDS” in **Table S2**); (3) a version with even stronger selection (no point-mass at zero and *α* = 1.07) to mimic the patterns at miRNAs (“miRNA” in **Table S2**); and (4) a version with substantially weaker selection (a 70% point-mass at zero and *α* = 0.55) to mimic the patterns at TFBSs (“TFBS” in **Table S2**).

When selecting the DFE from ref. [8], we chose the parameters estimated with a lower mutation rate (1.5 × 10^-8^), which was close to the one assumed for this study. In addition, when defining DFEs in terms of *s*_het_, we reduced the reported DFE by a scale factor of 4*N_e_* (using the estimated value of *N_e_*=12,378) to account for both that a population-scaled DFE was inferred in ref. [8] (accounting for a factor of 2*N_e_*) and that the inferred values of *s* are equivalent to 2shet under an additive model. This scaling was accomplished by reducing the value of *θ* in the inferred Gamma distribution from 820.6 to 0.0168. Notably, the mean of the DFE estimated for the 1000 Genomes Project data was intermediate between those estimated for the ESP European and LuCAMP data sets in ref. [8].

In each case, we simulated data with the assumed DFE for new mutations, denoted *f*(*x*), and then traced the DFE for the rare variants that remained in each data set after selection had been applied, denoted *g*(*x*). We then could estimate the DFE for the missing rare variants measured by ExtRaINSIGHT as 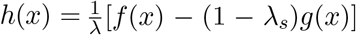, assuming that the full DFE can be expressed as a mixture of *g*(*x*) with weight 1 – *λ_s_* and *h*(*x*) with weight *λ_s_*. This mixture must also account for common variants, but we omit them because they occur at only a small fraction of sites in our setting.

## Data Availability

ExtRaINSIGHT and INSIGHT2 scores can be computed for any user-defined set of annotations using the ExtRaINSIGHT web portal at http://compgen.cshl.edu/extrainsight. The source code for the ExtRaINSIGHT server is available at https://github.com/CshlSiepelLab/extraINSIGHT.

## Acknowledgments

We thank Dr. Daniel Balick for providing simulation code from reference [18]. This research was supported by US National Institutes of Health grant R35-GM127070 and the Simons Center for Quantitative Biology at Cold Spring Harbor Laboratory. The content is solely the responsibility of the authors and does not necessarily represent the official views of the US National Institutes of Health.

## References

[1] Haldane JBS. The effect of variation of fitness. The American Naturalist. 1937;71:337–349.

[2] Fisher RA. On the dominance ratio. Proceedings of the Royal Society of Edinburgh. 1922;42:321–341.

[3] Haldane JBS. A mathematical theory of natural and artificial selection, part V: selection and mutation. In: Mathematical Proceedings of the Cambridge Philosophical Society. vol. 23. Cambridge University Press; 1927. p. 838–844.

[4] Eyre-Walker A, Keightley PD. The distribution of fitness effects of new mutations. Nat Rev Genet. 2007;8(8):610–618.

[5] Bataillon T, Bailey SF. Effects of new mutations on fitness: insights from models and data. Ann N Y Acad Sci. 2014;1320:76–92.

[6] Eyre-Walker A, Woolfit M, Phelps T. The distribution of fitness effects of new deleterious amino acid mutations in humans. Genetics. 2006;173(2):891–900.

[7] Boyko AR, Williamson SH, Indap AR, Degenhardt JD, Hernandez RD, Lohmueller KE, et al. Assessing the evolutionary impact of amino acid mutations in the human genome. PLoS Genet. 2008;4(5):e1000083.

[8] Kim BY, Huber CD, Lohmueller KE. Inference of the Distribution of Selection Coefficients for New Nonsynonymous Mutations Using Large Samples. Genetics. 2017;206(1):345–361.

[9] Huang YF, Siepel A. Estimation of allele-specific fitness effects across human protein-coding sequences and implications for disease. Genome Res. 2019;29(8):1310–1321.

[10] Kondrashov AS. Contamination of the genome by very slightly deleterious mutations: why have we not died 100 times over? J Theor Biol. 1995;175(4):583–594.

[11] Cassa CA, Weghorn D, Balick DJ, Jordan DM, Nusinow D, Samocha KE, et al. Estimating the selective effects of heterozygous protein-truncating variants from human exome data. Nat Genet. 2017;49(5):806–810.

[12] Lek M, Karczewski KJ, Minikel EV, Samocha KE, Banks E, Fennell T, et al. Analysis of proteincoding genetic variation in 60,706 humans. Nature. 2016;536(7616):285–291.

[13] Karczewski KJ, Francioli LC, Tiao G, Cummings BB, Alföldi J, Wang Q, et al. The mutational constraint spectrum quantified from variation in 141,456 humans. Nature. 2020;581(7809):434–443.

[14] Petrovski S, Wang Q, Heinzen EL, Allen AS, Goldstein DB. Genic intolerance to functional variation and the interpretation of personal genomes. PLoS Genet. 2013;9(8):e1003709.

[15] Samocha KE, Robinson EB, Sanders SJ, Stevens C, Sabo A, McGrath LM, et al. A framework for the interpretation of de novo mutation in human disease. Nat Genet. 2014;46(9):944–950.

[16] Havrilla JM, Pedersen BS, Layer RM, Quinlan AR. A map of constrained coding regions in the human genome. Nat Genet. 2019;51(1):88–95.

[17] Fuller ZL, Berg JJ, Mostafavi H, Sella G, Przeworski M. Measuring intolerance to mutation in human genetics. Nat Genet. 2019;51(5):772–776.

[18] Weghorn D, Balick DJ, Cassa C, Kosmicki JA, Daly MJ, Beier DR, et al. Applicability of the Mutation-Selection Balance Model to Population Genetics of Heterozygous Protein-Truncating Variants in Humans. Mol Biol Evol. 2019;36(8):1701–1710.

[19] Gronau I, Arbiza L, Mohammed J, Siepel A. Inference of natural selection from interspersed genomic elements based on polymorphism and divergence. Mol Biol Evol. 2013;30(5):1159–1171.

[20] Arbiza L, Gronau I, Aksoy BA, Hubisz MJ, Gulko B, Keinan A, et al. Genome-wide inference of natural selection on human transcription factor binding sites. Nat Genet. 2013;45(7):723–729.

[21] Li WH, Gojobori T, Nei M. Pseudogenes as a paradigm of neutral evolution. Nature. 1981;292(5820):237–239.

[22] Kimura M. Rare variant alleles in the light of the neutral theory. Mol Biol Evol. 1983;1(1):84–93.

[23] Kondrashov AS, Crow JF. A molecular approach to estimating the human deleterious mutation rate. Hum Mutat. 1993;2(3):229–234.

[24] Siepel A, Bejerano G, Pedersen JS, Hinrichs AS, Hou M, Rosenbloom K, et al. Evolutionarily conserved elements in vertebrate, insect, worm, and yeast genomes. Genome Res. 2005;15(8):1034–1050.

[25] Cooper GM, Stone EA, Asimenos G, Green ED, Batzoglou S, Sidow A. Distribution and intensity of constraint in mammalian genomic sequence. Genome Res. 2005;15(7):901–913.

[26] Gaffney DJ, Blekhman R, Majewski J. Selective constraints in experimentally defined primate regulatory regions. PLoS Genet. 2008;4(8):e1000157.

[27] Frankish A, Diekhans M, Ferreira AM, Johnson R, Jungreis I, Loveland J, et al. GENCODE reference annotation for the human and mouse genomes. Nucleic Acids Research. 2018;47(D1):D766–D773. doi:10.1093/nar/gky955.

[28] Lynch M. Rate, molecular spectrum, and consequences of human mutation. Proc Natl Acad Sci U S A. 2010;107(3):961–968.

[29] Fabregat A, Sidiropoulos K, Garapati P, Gillespie M, Hausmann K, Haw R, et al. The Reactome pathway Knowledgebase. Nucleic Acids Res. 2016;44(D1):D481–487.

[30] Bejerano G, Pheasant M, Makunin I, Stephen S, Kent WJ, Mattick JS, et al. Ultraconserved elements in the human genome. Science. 2004;304(5675):1321–1325.

[31] Pollard KS, Salama SR, Lambert N, Lambot MA, Coppens S, Pedersen JS, et al. An RNA gene expressed during cortical development evolved rapidly in humans. Nature. 2006;443(7108):167–172.

[32] Pollard KS, Salama SR, King B, Kern AD, Dreszer T, Katzman S, et al. Forces shaping the fastest evolving regions in the human genome. PLoS Genet. 2006;2(10):e168.

[33] Kostka D, Hubisz MJ, Siepel A, Pollard KS. The role of GC-biased gene conversion in shaping the fastest evolving regions of the human genome. Mol Biol Evol. 2012;29(3):1047–1057.

[34] Bejerano G, Lowe CB, Ahituv N, King B, Siepel A, Salama SR, et al. A distal enhancer and an ultraconserved exon are derived from a novel retroposon. Nature. 2006;441(7089):87–90.

[35] Prabhakar S, Visel A, Akiyama JA, Shoukry M, Lewis KD, Holt A, et al. Human-specific gain of function in a developmental enhancer. Science. 2008;321(5894):1346–1350.

[36] Scally A. The mutation rate in human evolution and demographic inference. Curr Opin Genet Dev. 2016;41:36–43.

[37] Ernst J, Kheradpour P, Mikkelsen TS, Shoresh N, Ward LD, Epstein CB, et al. Mapping and analysis of chromatin state dynamics in nine human cell types. Nature. 2011;473(7345):43–49.

[38] Hoffman MM, Ernst J, Wilder SP, Kundaje A, Harris RS, Libbrecht M, et al. Integrative annotation of chromatin elements from ENCODE data. Nucleic Acids Res. 2013;41(2):827–841.

[39] Kundaje A, Meuleman W, Ernst J, Bilenky M, Yen A, Heravi-Moussavi A, et al. Integrative analysis of 111 reference human epigenomes. Nature. 2015;518(7539):317–330.

[40] Sabarinathan R, Mularoni L, Deu-Pons J, Gonzalez-Perez A, López-Bigas N. Nucleotide excision repair is impaired by binding of transcription factors to DNA. Nature. 2016;532:264–267.

[41] Frigola J, Sabarinathan R, Gonzalez-Perez A, Lopez-Bigas N. Variable interplay of UV-induced DNA damage and repair at transcription factor binding sites. Nucleic Acids Research. 2020;49:891–901.

[42] Zerbino DR, Wilder SP, Johnson N, Juettemann T, Flicek PR. The Ensembl regulatory build. Genome Biol. 2015;16:56.

[43] Katzman S, Kern AD, Bejerano G, Fewell G, Fulton L, Wilson RK, et al. Human genome ultraconserved elements are ultraselected. Science. 2007;317(5840):915.

[44] Nóbrega MA, Zhu Y, Plajzer-Frick I, Afzal V, Rubin EM. Megabase deletions of gene deserts result in viable mice. Nature. 2004;431(7011):988–993.

[45] Snetkova V, Ypsilanti AR, Akiyama JA, Mannion BJ, Plajzer-Frick I, Novak CS, et al. Ultraconserved enhancer function does not require perfect sequence conservation. Nat Genet. 2021;53(4):521–528.

[46] Agrawal AF, Whitlock MC. Inferences about the distribution of dominance drawn from yeast gene knockout data. Genetics. 2011;187(2):553–566.

[47] Simmons MJ, Crow JF. Mutations affecting fitness in Drosophila populations. Annu Rev Genet. 1977;11:49–78.

[48] Kimura M, Ohta T. The average number of generations until extinction of an individual mutant gene in a finite population. Genetics. 1969;63(3):701–9.

[49] Rice WR. The high abortion cost of human reproduction. bioRxiv. 2018; p. 372193. doi:10.1101/372193.

[50] Wang X, Chen C, Wang L, Chen D, Guang W, French J. Conception, early pregnancy loss, and time to clinical pregnancy: a population-based prospective study. Fertil Steril. 2003;79(3):577–584.

[51] Torgerson DG, Boyko AR, Hernandez RD, Indap A, Hu X, White TJ, et al. Evolutionary processes acting on candidate cis-regulatory regions in humans inferred from patterns of polymorphism and divergence. PLoS Genet. 2009;5(8):e1000592.

[52] Kuhn RM, Haussler D, Kent WJ. The UCSC genome browser and associated tools. Briefings in bioinformatics. 2013;14(2):144–161.

[53] Gulko B, Hubisz MJ, Gronau I, Siepel A. A method for calculating probabilities of fitness consequences for point mutations across the human genome. Nat Genet. 2015;47(3):276–283.

[54] Gulko B, Siepel A. An evolutionary framework for measuring epigenomic information and estimating cell-type-specific fitness consequences. Nat Genet. 2019;51(2):335–342.

[55] Doan RN, Bae BI, Cubelos B, Chang C, Hossain AA, Al-Saad S, et al. Mutations in Human Accelerated Regions Disrupt Cognition and Social Behavior. Cell. 2016;167:341–354.e12.

[56] Dimitrieva S, Bucher P. UCNEbase—a database of ultraconserved non-coding elements and genomic regulatory blocks. Nucleic Acids Research. 2012;41(D1):D101–D109.

[57] Yang RY, Quan J, Sodaei R, Aguet F, Segrè AV, Allen JA, et al. A systematic survey of human tissuespecific gene expression and splicing reveals new opportunities for therapeutic target identification and evaluation. bioRxiv. 2018;doi:10.1101/311563.

