## Supplement for "Extreme purifying selection against point mutations in the human genome"

### Supplemental Material for: Extreme purifying selection against point mutations in the human genome

#### Supplemental Text

##### Approximation of the expected number of generations until extinction as a fraction of the neutral expectation

By Kimura and Ohta's formulas [1], the expected number of generations until a new mutant (at initial frequency $\frac{1}{2N}$) is lost from a finite population, as a function of the current population size $N$ and the scaled selection coefficient $S = 2N_e s$, where $N_e$ is the effective population size, is approximately given by,

$$t(S) = \frac{2N_e}{N}\left[\ln\left(\frac{N}{S}\right) + 1 - \gamma\right], \qquad \gamma = 0.577\ldots, \tag{1}$$

in the case of a semidominant deleterious mutation (with $h = \frac{1}{2}$), and by,

$$t(0) = \frac{2N_e}{N}\ln(2N) \tag{2}$$

in the case of a neutral mutation. In the regime of interest, $\ln(N/S) \gg 1$, so the ratio of these quantities can be roughly approximated as,

$$\frac{t(S)}{t(0)} \approx \frac{\ln(N) - \ln(S)}{\ln(2N)} = \frac{\ln(N) - \ln(S)}{\ln(N) + \ln 2}. \tag{3}$$

As discussed in the text, we estimate that ultraselected sites have values of $s_{\text{het}} = \frac{1}{2}s$ of about 0.03. Assuming the typical value of $N_e = 10^4$ for human populations, $S = 2N_e s = 4N_e s_{\text{het}} = 1200$, meaning that $\ln(S) \approx 7.1$. It is more difficult to know what $N$ should be in this setting, but Weghorn et al. [2] have argued for a plausible range of 0.5–8.0$\times 10^6$ based on current demographic models for human populations. Thus, $\ln(N)$ ranges from about 13.1 to 15.9, and we obtain values for $\frac{t(S)}{t(0)}$ ranging from 0.43 to 0.53.

### References

[1] Kimura M, Ohta T. The average number of generations until extinction of an individual mutant gene in a finite population. Genetics. 1969;63(3):701–9.

[2] Weghorn D, Balick DJ, Cassa C, Kosmicki JA, Daly MJ, Beier DR, et al. Applicability of the Mutation-Selection Balance Model to Population Genetics of Heterozygous Protein-Truncating Variants in Humans. Mol Biol Evol. 2019;36(8):1701–1710.

[3] Yang RY, Quan J, Sodaei R, Aguet F, Segrè AV, Allen JA, et al. A systematic survey of human tissue-specific gene expression and splicing reveals new opportunities for therapeutic target identification and evaluation. bioRxiv. 2018;doi:10.1101/311563.

[4] Kim BY, Huber CD, Lohmueller KE. Inference of the Distribution of Selection Coefficients for New Nonsynonymous Mutations Using Large Samples. Genetics. 2017;206(1):345–361.

Table S1: Ultraselection across the human genome (less conservative estimates)

| Feature | $\lambda_s$ | $\pm$ (stderr) | no. sites (M) | prop. sites | exp no. (M)[a] | exp. prop.[b] | fold enrich. | exp. lethal[c] | $s_{\text{het}}$ |
|---|---|---|---|---|---|---|---|---|---|
| CDS | 0.149 | 0.002 | 33.8 | 1.18% | 4.9 | 31.6% | 26.8 | 0.12 | - |
| $5'$ UTR | $-0.158$ | 0.002 | 8.2 | 0.29% | 0.0 | 0.0% | 0.0 | 0.00 | - |
| $3'$ UTR | 0.023 | 0.002 | 36.1 | 1.26% | 0.7 | 4.6% | 3.6 | 0.02 | - |
| splice | 0.464 | 0.002 | 0.8 | 0.03% | 0.4 | 2.3% | 85.0 | 0.01 | 2.0% |
| nonconserved lncRNA[d] | 0.008 | 0.002 | 453.6 | 15.78% | 1.8 | 11.8% | 0.7 | 0.04 | - |
| conserved lncRNA[e] | 0.055 | 0.002 | 23.3 | 0.81% | 1.2 | 7.7% | 9.5 | 0.03 | - |
| nonconserved intron[d] | 0.008 | 0.002 | 972.6 | 33.83% | 4.2 | 26.8% | 0.8 | 0.10 | - |
| conserved intron[e] | 0.057 | 0.002 | 44.3 | 1.54% | 2.4 | 15.3% | 9.9 | 0.06 | - |
| nonconserved intergenic[d] | 0.003 | 0.002 | 1255.5 | 43.67% | 0.0 | 0.0% | 0.0 | 0.00 | - |
| conserved intergenic[e] | 0.051 | 0.002 | 46.9 | 1.63% | 2.2 | 14.2% | 8.7 | 0.05 | - |
| Total | | | 2875.1 | 100.00% | 15.6 | 100.0% | | 0.43 | |

[a] Expected number of ultraselected sites after adjusting for background. In this case, the estimate for nonconserved intergenic regions (0.003) was subtracted from each estimate of $\lambda_s$ (see **Table 1** for a more conservative correction).

[b] Expected proportion of ultraselected sites after adjusting for background.

[c] Expected number of new lethal or near-lethal mutations per diploid individual, assuming a mutation rate of $1.2 \times 10^{-8}$ per generation per site.

[d] Sites not classified as conserved by phastCons.

[e] Sites classified as conserved by phastCons.

| Distribution | $\alpha^a$ | $\theta^a$ | ${\pi_0}^b$ | mean $g(x)$ | mean $f(x)$ | mean $h(x)$ | $\lambda_s$ | estimated $s_{\text{het}}$ |
|---|---|---|---|---|---|---|---|---|
| Kim et al., | 0.1930 | 0.0168 | 3.1% | 0.0023 | 0.0032 | 0.0303 | 0.0416 | - |
| 0d CDS | 0.8678 | 0.0168 | 3.1% | 0.0101 | 0.0141 | 0.0275 | 0.2340 | 0.0242 |
| miRNA | 1.0700 | 0.0168 | 0.0% | 0.0137 | 0.0189 | 0.0312 | 0.3396 | 0.0316 |
| TFBS | 0.5500 | 0.0168 | 70.0% | 0.0017 | 0.0028 | 0.0277 | 0.0275 | - |

[a]Parameters of assumed Gamma distribution, where $\alpha$ is the shape parameter and $\theta$ is the scale parameter
[b]Weight of point mass at zero.

Table S2: Means of full simulated DFE ($f(x)$), DFE associated with remaining rare variants ($g(x)$), and DFE inferred to be associated with the "missing" rare variants ($h(x)$) by mixture decomposition (see **Methods**). Also shown are the estimated values of $\lambda_s$ from simulated data, as well as the corresponding value of $s_{\text{het}}$ (for $\lambda_s > 0.18$).

### Supplemental Figures

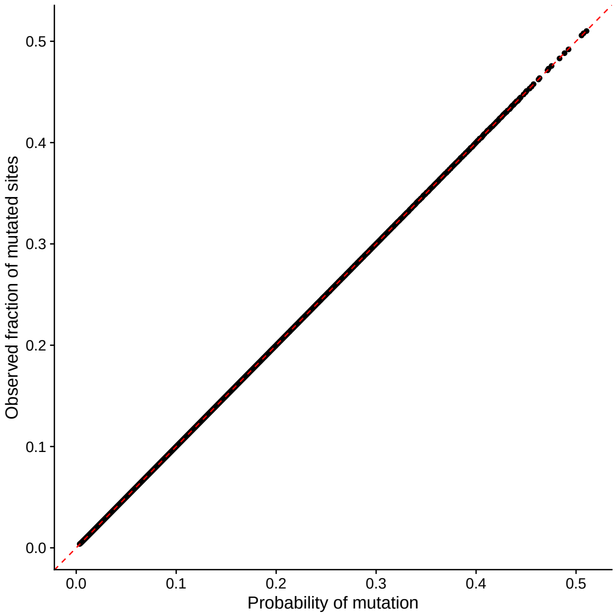


Figure S1: **Predicted vs. observed rates of rare variants in designated neutral regions.** Each point represents a single 50kb bin. Along the $x$-axis are the average values of $P_i$ across that bin, as predicted by our mutation model, and along the $y$ axis are the observed rates at which rare variants occur within that bin. The plot shows that the mutation model is well calibrated genome-wide for neutral sites.

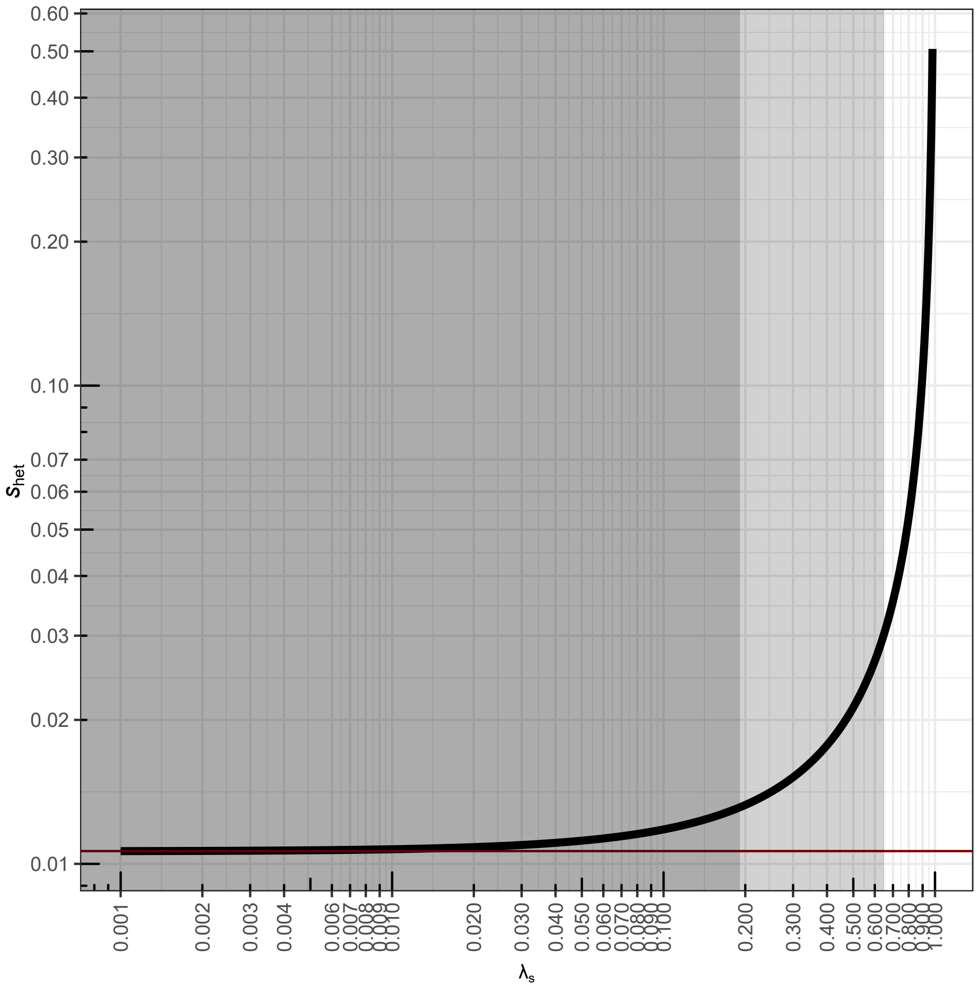


Figure S2: **Theoretical relationship between $\lambda_s$ and the selection coefficient against heterozygous mutations, $s_{\text{het}}$.** Curve represents equation 2 with $N = 71,702$ and $c = 1.35 \times 10^7$ based on our real data set (see **Methods**). The dark shaded region ($\lambda_s < 0.18$, $s_{\text{het}} < 0.013$) indicates the approximate regime where the relationship no longer yields an accurate estimator for $s_{\text{het}}$ with our data, and the lighter shaded region ($0.18 < \lambda_s < 0.65$, $0.013 < s_{\text{het}} < 0.03$) indicates the regime where the estimator is slightly inflated but still useful as a guide (see **Supplemental Fig. S3**).

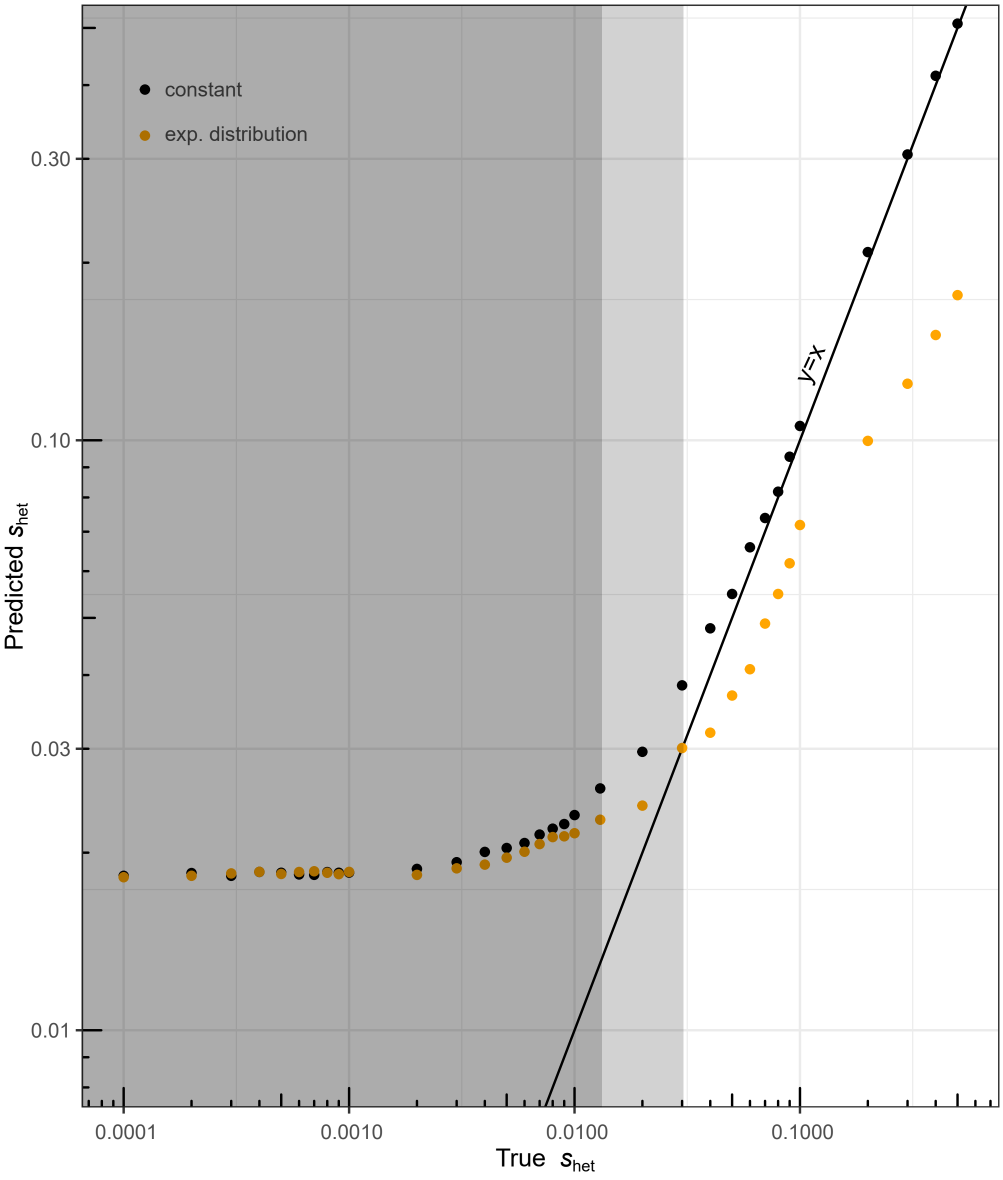


Figure S3: **True vs. predicted values of $s_{\text{het}}$ in simulation.** Data sets of 71,702 diploid individuals and 100,000 sites were simulated using software from ref. [2] with mean $s_{\text{het}}$ ranging from 0.0001 to 0.5 ($x$-axis). In one version, all sites were assigned the same "true" value of $s_{\text{het}}$ ("constant"; black points) and, in another, sitewise values of $s_{\text{het}}$ were drawn from an exponential distribution with the given mean value ("exp. distribution"; orange points). ExtRaINSIGHT was applied to each simulated data set, and then the estimated value of $\lambda_s$ was converted to a predicted $s_{\text{het}}$ ($y$-axis) using equation 2. All simulations assumed a European demographic history (see **Methods**). As in **Supplemental Fig. S2**, the dark and light gray regions respectively indicate the regimes in which the estimator for $s_{\text{het}}$ is no longer useful, and is inflated but still approximately useful.

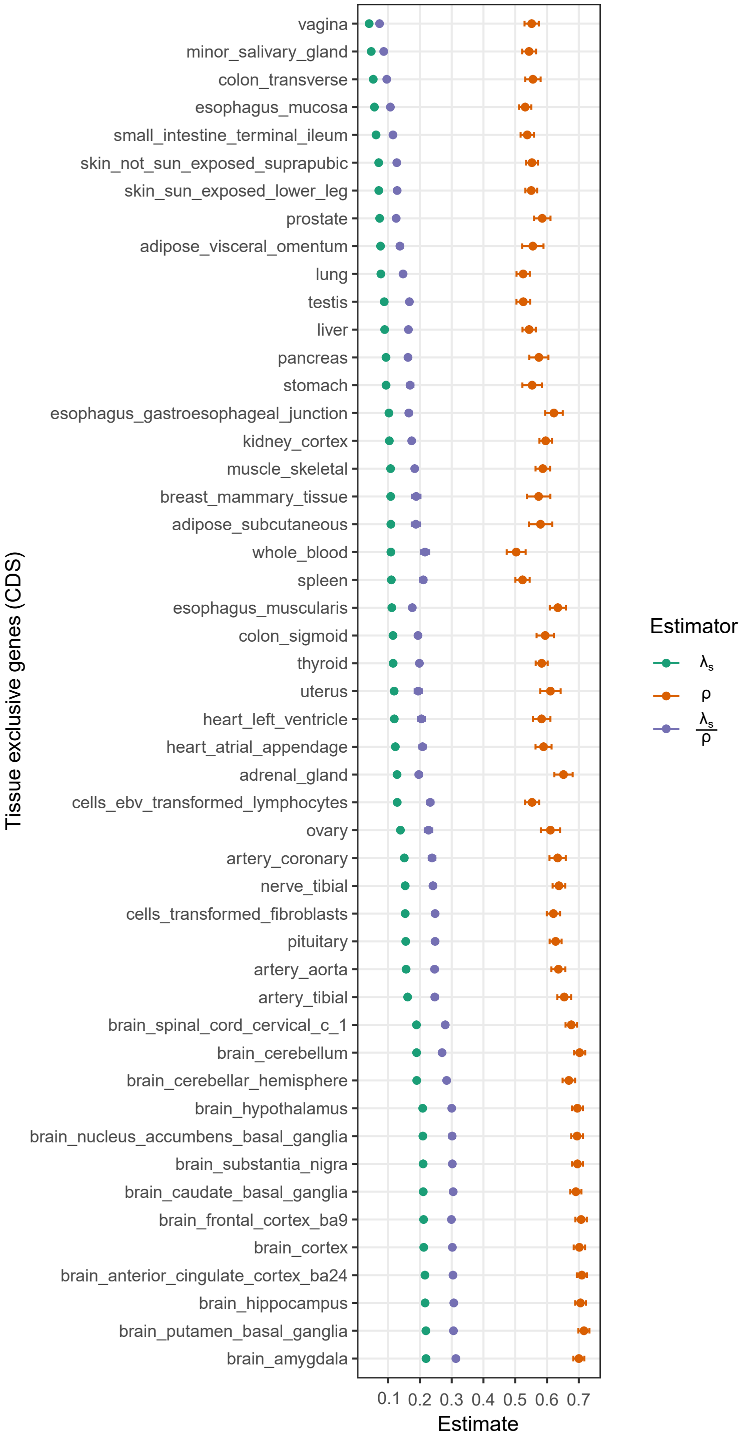


Figure S4: **Measures of purifying selection in protein-coding genes exhibiting tissue-specific gene expression.** Tissue-specific genes were obtained from ref. [3] as detailed in the **Methods** section. An estimate for each tissue is shown for both ExtRaINSIGHT ($\lambda_s$) and INSIGHT ($\rho$). Error bars indicate one standard error (see **Methods**).

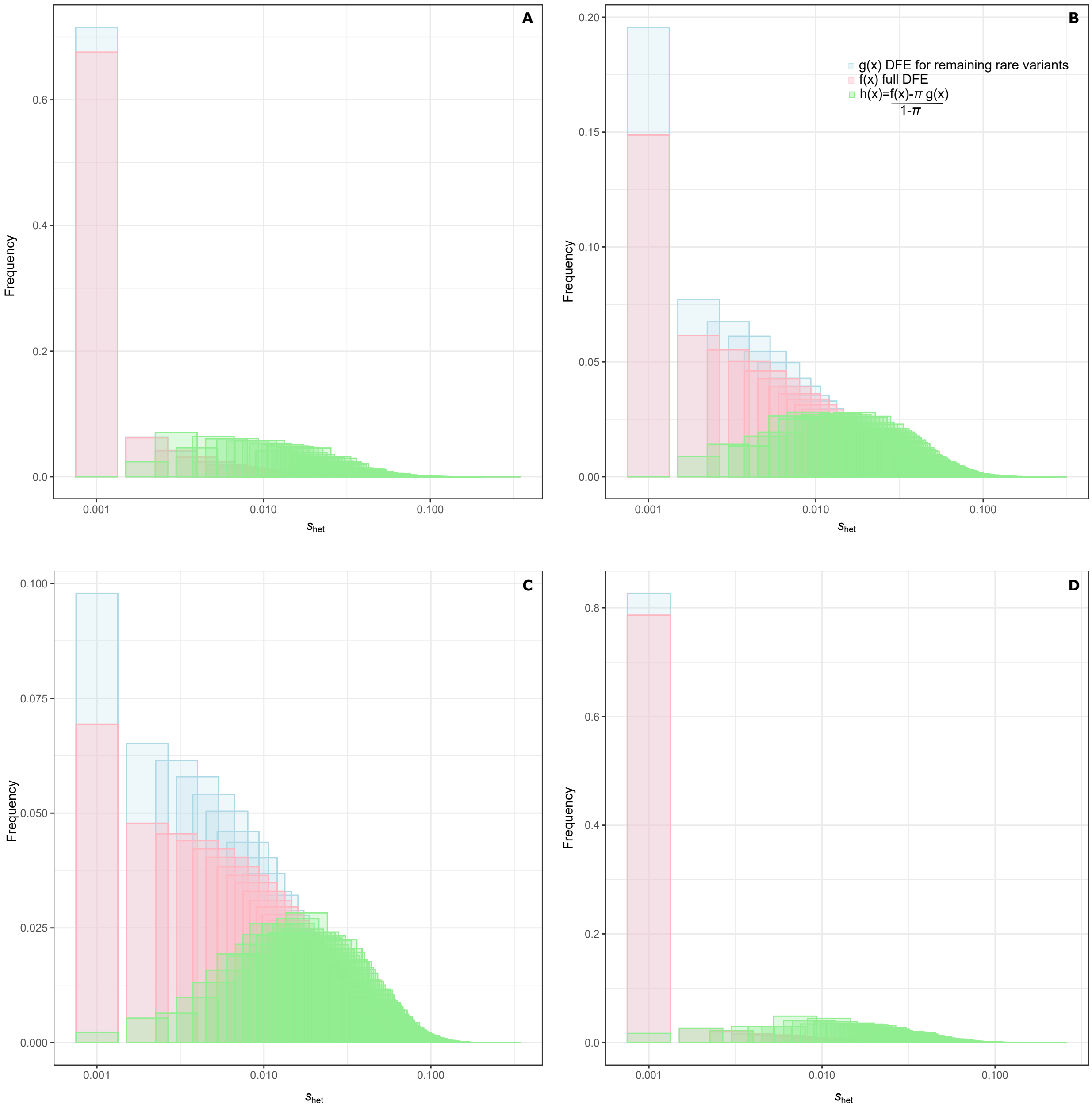


Figure S5: **Comparison of DFEs for all sites, rare variants that remain, and "missing" rare variants in simulations.** Simulated DFEs ($f(x)$; pink), DFEs for rare variants that remain in the data ($g(x)$; blue), and DFEs inferred by mixture decomposition for the rare variants that are missing ($h(x)$; green). Results are shown for four distinct DFEs: **(A)** a DFE published by Kim et al. [4] consisting of a mixture of a point-mass at zero (with weight 0.031) and a Gamma distribution with $\alpha$=0.1930 and $\theta$=0.0168. **(B)** a modified DFE designed to approximately match our observations at 0d sites in coding regions, consisting of a mixture of a point-mass at zero (weight 0.031) and a Gamma distribution with $\alpha$=0.8687 and $\theta$=0.0168. **(C)** a modified DFE designed to approximately match our observations at evolutionarily ancient miRNAs, equal to a Gamma distribution with $\alpha$=1.07 and $\theta$=0.0168. **(D)** a modified DFE designed to approximately match our observations at TFBS, consisting of a mixture of a point-mass at zero (with weight 70%) and a Gamma distribution with $\alpha$=0.55 and $\theta$=0.0168. Means of these distributions along with our $\lambda_s$ and $s_{\text{het}}$ estimates are shown in **Supplemental Table S2.**
